# Modulation of DNA polymerase IV activity by UmuD and RecA* observed by single-molecule time-lapse microscopy

**DOI:** 10.1101/620195

**Authors:** Sarah S. Henrikus, Amy E. McGrath, Slobodan Jergic, Matthew L. Ritger, Phuong T. Pham, Elizabeth A. Wood, Myron F. Goodman, Michael M. Cox, Antoine M. van Oijen, Harshad Ghodke, Andrew Robinson

**Author notes:** Corresponding author. Mailing address: School of Chemistry and Molecular Bioscience, University of Wollongong, Wollongong, NSW 2522, Australia.

## Abstract

DNA polymerase IV (pol IV) is expressed at increased levels in *Escherichia coli* cells suffering high levels of DNA damage. In a recent single-molecule imaging study, we demonstrated that elevating the pol IV concentration is not sufficient to provide access to binding sites on the nucleoid, suggesting that other factors may recruit pol IV to its substrates once the DNA becomes damaged. Here we extend this work, investigating the proteins UmuD and RecA as potential modulators of pol IV activity. UmuD promotes long-lived association of pol IV with the nucleoid, whereas its cleaved form, UmuD’, which accumulates in DNA-damaged cells, inhibits binding. In agreement with proposed roles for pol IV in homologous recombination, up to 40% of pol IV foci colocalise with a probe for RecA* nucleoprotein filaments in ciprofloxacin-treated cells. A hyperactive RecA mutant, *recA*(E38K), allows pol IV to bind the nucleoid even in the absence of exogenous DNA damage. *In vitro,* RecA(E38K) forms RecA*-like structures that can recruit pol IV, even on double-stranded DNA, consistent with a physical interaction between RecA and pol IV. Together, the results indicate that UmuD and RecA modulate the binding of pol IV to its DNA substrates, which frequently coincide with RecA* structures.

## Introduction

DNA polymerase IV (pol IV), encoded by *dinB*, is one of three specialised DNA polymerases that are produced at increased levels in *Escherichia coli* cells suffering DNA damage (Napolitano *et al*., 2000). *In vitro*, DNA polymerase IV is capable of translesion synthesis (TLS) on a variety of different lesion-containing DNA substrates (Kim *et al*., 1997; Jarosz *et al*., 2006; Kumari *et al*., 2008; Yuan *et al*., 2008; Cafarelli *et al*., 2014; Ikeda *et al*., 2014; Fuchs, 2016; Henrikus *et al*., 2018a). The most commonly proposed function for pol IV within cells is TLS at stalled replication forks, which may help to maintain chromosomal replication in cells experiencing DNA damage (Fuchs and Fujii, 2013; Kath *et al*., 2014). However, in the cell, the majority of binding sites for pol IV on the nucleoid appear distal to replisome markers (Henrikus *et al*., 2018b). There is significant evidence that pol IV participates in other pathways, including recombinational repair (Lovett, 2006; Williams *et al*., 2010; Shee *et al*., 2011; Shee *et al*., 2012b; Shee *et al*., 2012a; Pomerantz *et al*., 2013a; Pomerantz *et al*., 2013b) and transcription-coupled TLS (Cohen *et al*., 2009; Cohen *et al*., 2010; Kang *et al*., 2010; Cohen and Walker, 2011).

Single-molecule time-lapse imaging of fluorescently tagged pol IV in live *Escherichia coli* cells revealed that various DNA-damaging agents (ciprofloxacin, UV light and methyl methanesulfonate [MMS]) up-regulate the production of pol IV and create binding sites for pol IV on the nucleoid (Henrikus *et al*., 2018b). Only 10% of the pol IV binding events (pol IV foci) occurred in the vicinity of replisomes. At late time points during the SOS response (90–100 min after damage induction) pol IV continued to form foci but no longer colocalised with replisomes, even at low levels. This led to the hypothesis that replisome access might be controlled by protein–protein interactions that change around 90–100 min after the induction of SOS. The results also suggest that pol IV function is focused primarily on events that occur away from the replication fork. The recruitment of pol IV to the processivity factor β strongly depends on the source of DNA damage (Thrall *et al*., 2017), indicating that the type of DNA lesion and changes in metabolism may affect which repair pathway(s) pol IV participates in (Henrikus *et al*., 2018a).

The UmuD protein and its cleaved form UmuD′ have a potential role in regulating pol IV activity in cells (Godoy *et al*., 2007). The auto-cleavage of UmuD to the shorter form UmuD′ is induced by the cellular recombinase RecA, in particular RecA nucleoprotein filaments (denoted RecA*). UmuD cleavage (Burckhardt *et al*., 1988; Woodgate and Ennis, 1991; Frank *et al*., 1996) has long been understood to be a key step in the activation of the highly mutagenic enzyme DNA polymerase V (pol V) Mut (UmuD′_2_C-RecA-ATP; (Jiang *et al*., 2009)). Several lines of evidence suggest that the conversion of UmuD to UmuD′ might also regulate the activity of pol IV in *E. coli* (Godoy *et al*., 2007). Far-Western blots and co-purification experiments indicate that pol IV interacts with UmuD_2_ and UmuD′_2_, but not the heterodimer UmuDD′ (Godoy *et al*., 2007). Overexpression of pol IV induces high rates of –1 frameshift mutations in cells, that can be supressed by co-overexpression of UmuD, but not co-overexpression of UmuD′ (Godoy *et al*., 2007). Furthermore, UmuD and UmuD′ overexpression reduced frequencies in an adaptive mutagenesis assay compared to an empty vector; overproduction of UmuD even lowered frequencies to equivalent levels of a catalytically dead *dinB* mutant (Godoy *et al*., 2007). These observations have led to the proposal that UmuD status regulates the mutagenic activity of pol IV-dependent DNA synthesis. Despite these advances, it remains unclear if UmuD or UmuD′ solely affects the fidelity of pol IV, or if UmuD and UmuD′ might also regulate the DNA-binding activity of pol IV as a means to modulate pol IV-dependent mutagenesis.

A series of live-cell studies indicate that pol IV operates in the repair of double-strand breaks (DSBs) (Ponder *et al*., 2005; Foster, 2007; Shee *et al*., 2011; Rosenberg *et al*., 2012; Shee *et al*., 2012b; Mallik *et al*., 2015; Moore *et al*., 2017). Reducing DSB formation (by mitigating the destructive effects of reactive oxygen species) or introducing defects in the end-resection of double-strand breaks (Δ*recB* mutation) greatly reduces the number of pol IV foci formed in cells treated with ciprofloxacin or trimethoprim (Henrikus *et al*., 2019b). At end-resected DSBs, RecA nucleoprotein filaments facilitate repair through homologous recombination (Cox, 2007), suggesting that pol IV should colocalise with RecA* structures in cells engaged in DSB repair. A series of observations support this notion. Pol IV forms a physical interaction with RecA *in vitro* and this interaction modulates the fidelity of pol IV-dependent DNA synthesis (Godoy *et al*., 2007; Indiani *et al*., 2013; Cafarelli *et al*., 2014). This interaction is proposed to provide pol IV with the ability to participate in DNA synthesis during RecA-dependent strand exchange reactions (Tashjian *et al*., 2019). In a fluorescence microscopy study (Mallik *et al*., 2015), pol IV was shown to colocalise with RecA structures *in vivo*. However, the RecA-GFP probe that was used to observe RecA localisation does not differentiate between active forms of RecA (i.e. RecA*) and inactive forms, such as storage structures (Ghodke *et al*., 2019). Furthermore, this RecA-GFP (*recA4155-gfp*) probe is deficient in recombination, SOS induction and UV survival (Renzette *et al*., 2005). It therefore remains unclear whether RecA* structures, such as those that form as intermediates of recombination, represent major or minor substrates for pol IV in cells. With the recent development of a RecA*-specific probe, PAmCherry-mCI (Ghodke *et al*., 2019), we are now in a position to measure pol IV–RecA* colocalisation directly in a time-resolved manner.

In this work, we set out to test the following: 1. whether the UmuD cleavage status affects the extent of pol IV focus formation and pol IV colocalisation with a replisome marker and/or the lifetimes of pol IV molecules binding to its substrates, and 2. whether pol IV predominantly binds at RecA* structures. We use the drug ciprofloxacin, a DNA gyrase inhibitor, that induces DSBs upon treatment (Zhao *et al*., 1997). Using single-molecule live-cell imaging, we demonstrated that the binding of pol IV to the nucleoid is promoted by full-length UmuD in cells treated with the DNA damaging antibiotic ciprofloxacin. In contrast, UmuD′ diminishes pol IV binding. We observed that a large proportion of pol IV foci (up to 40%) colocalise with a RecA* marker in ciprofloxacin-treated cells. The *recA*(E38K) mutation (also known as *recA730*), which constitutively produces RecA*-like activity (Cazaux *et al*., 1993; Wang *et al*., 1993; Ennis *et al*., 1995), promotes the binding activity of pol IV to the nucleoid, even in the absence of DNA damage. We further showed that pol IV physically interacts with RecA(E38K), which forms RecA*-like structures on single-stranded as well as double-stranded DNA, suggesting that pol IV might also associate with these RecA*-like structures in cells. These findings provide evidence for regulatory roles for both UmuD and RecA in modulating the binding activity of pol IV in *E. coli* cells. RecA* structures that likely mark sites of on-going DSB repair appear to serve as major binding sites for pol IV in live cells treated with ciprofloxacin.

## Results

### Deletion of *umuDC* increases pol IV-τ colocalisation

In a previous study, we carried out time-lapse measurements on *E. coli* cells treated with DNA damaging agents (Henrikus *et al*., 2018b). We found that the colocalisation of pol IV foci with replisome markers started at ∼10% prior to treatment, and dropped to < 5% (i.e. baseline levels) at a time-point 90–100 after the onset of treatment. In a separate study, we observed that pol V (UmuC-mKate2) enters the cytosol and forms foci on the nucleoid at this same 90 min time-point (Robinson *et al*., 2015). This spatial re-distribution of the UmuC-mKate2 marker required cleavage of UmuD to UmuD′. The similar timing of the changes in pol IV and pol V localisation, together with established links between pol IV activity and UmuD/UmuD′ status described above, led us to hypothesise that UmuD cleavage and/or formation of pol V at the 90 min time-point alters the colocalisation of pol IV with replisome markers.

To investigate the effect of pol V and/or its precursors (UmuD and UmuC) on the extent of pol IV focus formation and colocalisation with a replisome marker, we constructed two strains: i) *dinB-YPet dnaX-mKate2 umuDC^+^* (EAW643, (Henrikus *et al*., 2018b)) and ii) *dinB-YPet dnaX-mKate2* Δ*umuDC* (SSH007). The *dnaX-mKate2* allele encodes for a fluorescent fusion of the τ clamp loader protein, serving as a marker for the replisome, τ-mKate2. We previously showed that the fluorescent protein fusion of DinB-YPet is fully functional, yielding pol IV-dependent mutagenesis activity upon ciprofloxacin treatment in both *dnaX*^+^ and *dnaX-mKate2* cells (Henrikus *et al*., 2018b; Henrikus *et al*., 2019b).

Time-lapse movies were recorded for each strain after treatment with ciprofloxacin (30 ng mL^−1^). At t = 0 min, images of the DinB-YPet signal and τ-mKate2 signal (replisome marker) were recorded for untreated cells. Directly after t = 0, ciprofloxacin was introduced to the flow cell and a time-lapse was recorded over a period of 3 h. We previously showed that a catalytically dead mutant DinB(D103N)-YPet does not form foci under these imaging conditions (Henrikus *et al*., 2018b). This suggests that the DinB-YPet foci we normally detect are formed as DinB-YPet binds to the nucleoid and carries out DNA synthesis, at which point its diffusion is slowed sufficiently to produce a single-molecule focus. From the time-lapse movies, the numbers of DinB-YPet foci per cell, reflective of pol IV binding to the nucleoid (Henrikus *et al*., 2018b), were determined at 0, 30, 60, 90 and 120 min time points (Fig 1). Colocalisation between DinB-YPet foci and τ-mKate2 foci was also monitored. In order to enhance diffusional contrast in our images we used longer exposure times when capturing DinB-YPet signal (300 ms) than in our previous study (50 ms; (Henrikus *et al*., 2018b). We nonetheless recorded a complementary set of colocalisation measurements with the shorter exposure time of 50 ms in order to better capture transient foci and allow for more direct comparison with our previous results (**Fig S1**).

**Figure 1.**
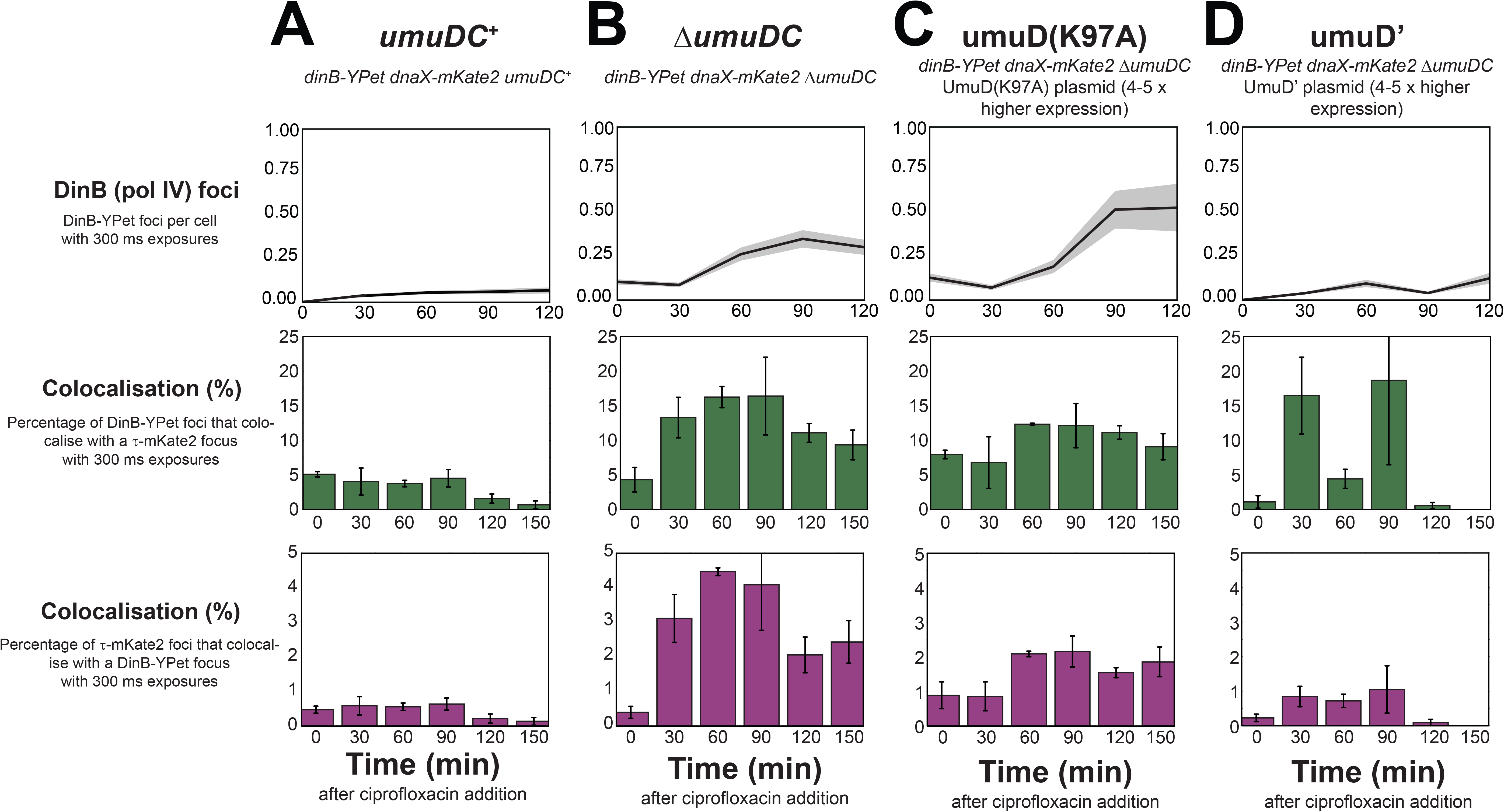
Number of DinB foci per cell and colocalisation measurements between DinB and τ in different *umuDC* mutants following ciprofloxacin treatment. (A) Upper panel: Number of DinB foci per cell in *umuDC*^+^ at 300 ms exposures. Error bar represents standard error of the mean for *n* > 100 cells. Middle panel: Colocalisation percentage of DinB with DnaX (green bars) in *umuDC*^+^. Time points are binned over 30 min. Error bar represents standard error of the mean between biological triplicates. Lower panel: Colocalisation percentage of DnaX with DinB (magenta bars) in *umuDC*^+^. Time points are binned over 30 min. The other columns represent the same measurements as in (A), except from the cell strains (B) Δ*umuDC*, (C) Δ*umuDC* + UmuD(K97A) expressed from a plasmid, and (D) Δ*umuDC* + UmuD’ expressed from a plasmid.

We first monitored pol IV behaviour in cells expressing wild-type levels of UmuD and UmuC (EAW643, Table 1). Cells exhibited very few pol IV foci prior to ciprofloxacin treatment (Fig 1A, upper panel), as observed previously (Henrikus *et al*., 2018b). After ciprofloxacin addition, the number of pol IV foci per cell increased to an average of 0.1 foci per cell from 60 min, i.e. one in ten cells exhibited a pol IV focus. Consistent with our previous observations (Henrikus *et al*., 2018b), the percentage of pol IV foci that colocalised with τ-mKate2 dropped markedly between the 90 min and 120 min time-points (Fig 1A, middle panel). From 0–90 min after ciprofloxacin addition, 5% of pol IV foci colocalised with the replisome marker τ. From 120–150 min this decreased to < 2%. These values are somewhat lower than those we reported previously (10% dropping to < 5%) and is attributable to the longer image exposure times used in the current study (**Fig S1**). The percentage of τ foci that contained a pol IV focus followed a similar trend (Fig 1A, lower panel); from 0–90 min after ciprofloxacin addition, 0.5% of τ foci contained a pol IV focus, dropping to ∼0.1% (indistinguishable from chance colocalisation) from 120–150 min.

**Table 1.** Strains used in this study.

| Strain | Relevant Genotype | Parent strain | Source/technique |
| --- | --- | --- | --- |
| MG1655 | *dinB*<sup>+</sup> *umuDC*<sup>+</sup> *lexA*<sup>+</sup> *recA*<sup>+</sup> | - | (Blattner *et al.*, 1997) |
| RW1594 | *dinB*-YPet *dnaX*-*mKate2*<br>*sulA*::*kan*<sup>R</sup> *lexA*(Def) Cm<sup>R</sup> | RW1588 | (Henrikus *et al.*, 2018b) |
| RW244 | *recA*(E38K) *srlD300*::Tn10 | - | (McDonald *et al.*, 1998) |
| RW1598 | *dinB*-YPet *dnaX*-*mKate2*<br>*sulA*::*kan*<sup>R</sup> *lexA*(Def)::Cm<sup>R</sup><br>*recA*(E38K) *srlD300*::Tn10 | RW1594 | Transduction of RW1594 with P1 grown on RW244 |
| EAW633 | *dinB*-YPet::kan<sup>R</sup> | MG1655 | (Henrikus *et al.*, 2018b) |
| EAW643 | *dinB*-YPet::FRT *dnaX*-<br>*mKate2*::kan<sup>R</sup> *lexA*<sup>+</sup> | EAW633 | (Henrikus *et al.*, 2018b) |
| RW880 | Δ*umuDC*::Cm<sup>R</sup> | MG1655 | (Henrikus *et al.*, 2019b) |
| SSH007 | *dinB*-YPet::FRT *dnaX*-<br>*mKate2*::kan<sup>R</sup> *lexA*<sup>+</sup><br>Δ*umuDC*::Cm<sup>R</sup> | EAW643 | Transduction of EAW643 with P1 grown on RW880 |
| SSH007 +<br>pJM1243 | *dinB*-YPet::FRT *dnaX*-<br>*mKate2*::kan<sup>R</sup> *lexA*<sup>+</sup><br>Δ*umuDC*::Cm<sup>R</sup> (chr) +<br>UmuD(K97A) (pl) | SSH007 | Transformation of SSH007 with pJM1243 |
| SSH007+<br>pRW66 | *dinB*-YPet::FRT *dnaX*-<br>*mKate2*::kan<sup>R</sup> *lexA*<sup>+</sup><br>Δ*umuDC*::Cm<sup>R</sup> (chr) +<br>UmuD' (pl) | SSH007 | Transformation of SSH007 with pRW66 |
| SSH092 | *dinB*-YPet::kan<sup>R</sup> (chr) +<br>PAmCherry-mCI (pl) | EAW633 | Transformation of EAW633 with pJMuvrA-PAmCherry-mCI (Ghodke *et al.*, 2019) |

We next examined the effect of deleting the *umuDC* operon (and thus eliminating UmuD and UmuC) on the number of pol IV foci and the extent of colocalisation with τ foci (SSH007, Table 1). From 30 min, 10–15% of pol IV foci colocalised with replisomes. Compared to *umuDC^+^* cells, Δ*umuDC* cells exhibited a three-fold increase in the number pol IV foci per cell with ∼0.3 foci per cell from 60 min after ciprofloxacin addition (Fig 1B, upper panel). Moreover, deletion of *umuDC* led to a three-fold increase in the percentage of pol IV foci that colocalise with a τ focus (Fig 1B, middle panel). Interestingly, pol IV-τ colocalisation now persisted above 10% for the 90, 120 and 150 min time points. The percentage of τ foci that contained a pol IV focus was also elevated in the Δ*umuDC* background (Fig 1B, lower panel). From 30 min, 2–4% of τ foci contained a pol IV focus. Compared to *umuDC^+^* cells, this represents a six-to eight-fold increase in colocalisation.

Taken together, the time-lapse imaging results show that cells lacking *umuDC* exhibit an increase in the number of pol IV foci per cell, accompanied by enhanced pol IV-τ colocalisation during the late SOS response (90–120 min). In cells lacking *umuDC*, the maximum extent of pol IV-τ colocalisation is 15%. This suggests that in cells lacking UmuD and UmuC, replisomes still do not represent the major binding substrate for pol IV.

### Cleavage state of UmuD affects the binding behaviour of pol IV

The increased numbers of pol IV foci and increased pol IV-τ colocalisation in Δ*umuDC* than in *umuDC*^+^ cells could manifest through two scenarios: 1. the deletion of the *umuDC* operon, which encodes for pol V, eliminates competition between pol IV and pol V for binding sites on the nucleoid. 2. a subunit of pol V has a regulatory effect on pol IV focus formation and pol IV-τ colocalisation. It has been shown previously that UmuD_2_ and UmuD′_2_ physically interact with pol IV and modulate its mutagenic activity (Godoy *et al*., 2007). To that end, we tested if UmuD or UmuD′ affect the extent of pol IV focus formation and the colocalisation between pol IV with τ, in the absence of UmuC (and thus pol V).

We constructed two strains, both of which include the *dinB-YPet* and *dnaX-mKate2* alleles: i) Δ*umuDC* (SSH007) expressing the non-cleavable UmuD(K97A) protein from a low-copy plasmid (SSH007 + pJM1243), and ii) SSH007 expressing the ‘cleaved’ UmuD′ protein from a low-copy plasmid (SSH007 + pRW66). The amount of UmuD(K97A) and UmuD′ produced from each plasmid is 4–5-fold higher than UmuD expressed from its native chromosomal locus (Churchward *et al*., 1984). Time-lapse analysis was repeated as described above.

We first explored the effects of expressing the non-cleavable UmuD(K97A) mutant in *dinB-YPet dnaX-mKate2* Δ*umuDC* cells (SSH007 + pJM1243, Table 1). At the 90 min time point, cells contained on average 0.6 pol IV foci per cell — a six-fold increase over *umuDC^+^* cells (Fig 1C, upper panel). This sixfold increase in pol IV foci per cell was accompanied by a three-fold increase in colocalisation with the replisome marker τ-mKate2 (Fig 1C, middle panel). From 30 min after damage induction, 13% of pol IV foci overlapped with a τ focus. This colocalisation remained relatively constant during the later stages of the SOS response; colocalisation did not drop below 9% from 90–120 min as observed in *umuDC^+^*cells. These observations reveal that UmuD(K97A), and by inference uncleaved UmuD, promote the binding of pol IV to DNA and do not limit pol IV-τ colocalisation beyond 90 min.

During the later stages of the SOS response (90 min after SOS induction), UmuD is cleaved to UmuD′ (Woodgate *et al*., 1989). To explore the effects of UmuD′ on pol IV behaviour, we imaged Δ*umuDC* cells expressing UmuD′ directly from a plasmid (SSH007 + pRW66, Table 1). These cells produced ∼0.1 DinB-YPet foci per cell at 60 min (Fig 1D, upper panel), similar to *umuDC^+^* cells. In the cells expressing UmuD′, colocalisation of pol IV with τ was generally low, but highly variable (Fig 1D, middle panel). Two large spikes in colocalisation were apparent at the 30 and 90 min time points. However, due to the low number of foci available for analysis at these time-points, there was very large error associated with these values. No spikes in colocalisation were observed when measuring the proportion of τ foci that contained a pol IV focus (Fig 1D, lower panel). Importantly, the colocalisation of pol IV with τ decreased to < 1% after 90 min (Fig 1D, middle panel). Similarly, the percentage of τ foci that contained a pol IV focus drops between the 90 and 120 min time points (Fig 1D, lower panel). From 30–90 min, ∼1% of τ foci contained a pol IV focus. By 120 min < 0.1% of τ foci contained a pol IV focus. Overall, the introduction of UmuD′ into Δ*umuDC* cells restores rates of focus formation and colocalisation with the replisome marker τ to near wild-type (*umuDC*^+^) levels.

Taken together, the time-lapse imaging results show that the presence of non-cleavable UmuD results in an increase in nucleoid binding by pol IV, accompanied by increased pol IV-τ colocalisation during the late SOS response (90–120 min). Strikingly, UmuD′ suppresses the formation of pol IV foci, also limiting pol IV-τ colocalisation. These results suggest that UmuD cleavage represents a biochemical switch that alters aspects of pol IV activity. Importantly, these effects were apparent in the absence of UmuC, ruling out the possibility that the drop in pol IV colocalization with replisomes at 90 min occurs because of competition for substrates between pols IV and V.

### UmuD(K97A) but not UmuD’ promotes long-lived pol IV binding events

Time-lapse imaging revealed differences in pol IV activity in *umuDC* variants with respect to the number of foci per cell and pol IV-τ colocalisation. We noted that in the various DinB-YPet images the foci formed in different strains appeared to exhibit differences in both intensity and shape (Figs 2A–D, first row; 300 ms exposures). For the *umuDC*^+^ (Fig 2A), Δ*umuDC* (Fig 2B) and UmuD′-expressing cells (Fig 2D), most foci were relatively faint and diffuse. In contrast, cells expressing UmuD(K97A) produced brighter, and much more distinct, pol IV foci. Reasoning that these differences might reflect differences in the nature of pol IV interactions with the substrates, we next measured the binding lifetime of pol IV at these sites. Image sets were recorded during three periods following the addition of ciprofloxacin: 20–45 min, 55–85 min and 120–180 min. For each time interval and each strain (EAW643, SSH007, SSH007 + pJM1243, SSH007 + pRW66; Table 1), burst acquisitions of the DinB-YPet signal were recorded (300 images of 34 ms exposure time, total length of 10.2 s). Subsequently, a corresponding image of the replisome marker τ-mKate2 was collected (see **Fig S2B** for imaging sequence).

**Figure 2.**
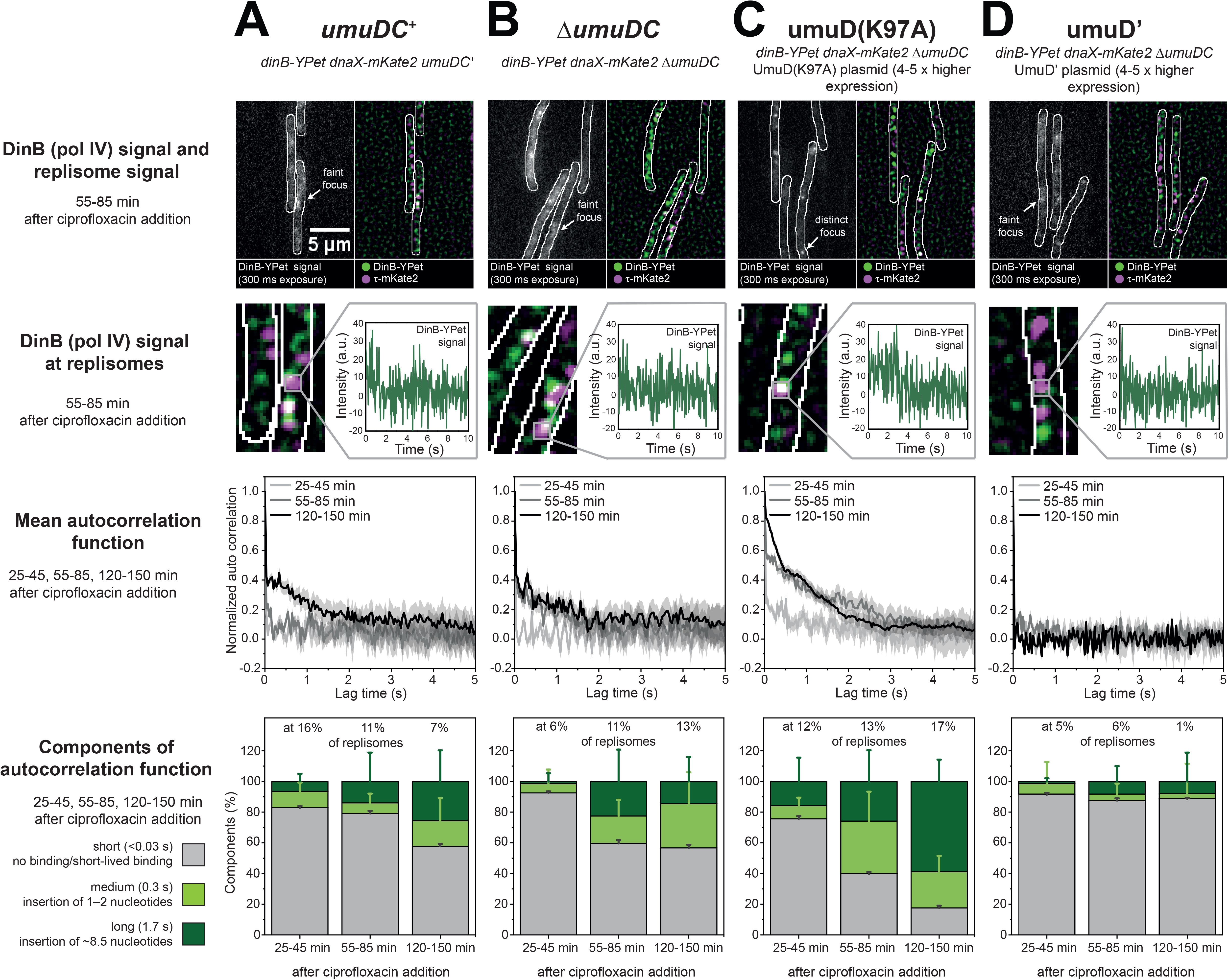
Binding activity of DinB at and away from replisomes in different *umuDC* mutants. (A) **Upper panel**: Images of DinB and DnaX signal in *umuDC*^+^. Left: Projection of DinB signal consistent with 300 ms exposure times. Right: Merged images of discoidal filtered DinB (green) and DnaX signal (magenta). **Second panel from the top**: Exemplary trajectory showing DinB activity at replisomes in *umuDC*^+^. **Third panel from the top**: Mean autocorrelation function showing DinB activity at replisomes in *umuDC*^+^ at 25–45 (light grey line), 55–85 (grey line) and 120–150 min (black line). Error bars represent standard error of the mean over > 100 trajectories. **Bottom panel**: Components of the autocorrelation function for DinB at replisomes in *umuDC*^+^ showing short (< 0.03 s, grey), medium (0.3 s, light green) and long components (1.7 s, dark green). The error bars for long and medium components were extracted from the fit error using the two-exponential fit (**Suppl.** Fig 1G, H). The error bar from the short components is equivalent to the standard error of the mean from the mean autocorrelation function at lag time 0s. (B) similar to (A), however in Δ*umuDC*. (C) similar to (A), however in Δ*umuDC* + UmuD(K97A) expressed from a plasmid. (D) similar to (A), however in Δ*umuDC* + UmuD’ expressed from a plasmid.

For the *umuDC*^+^, Δ*umuDC* and UmuD′-expressing cells, intensity trajectories collected at the positions of τ foci predominantly exhibited short-lived binding events (Fig 2, second row). Cells expressing UmuD(K97A), on the other hand, often produced long-lived pol IV binding events. To comprehensively assess the binding lifetimes of pol IV with respect to the UmuD status at sites of the replisome marker, mean autocorrelation functions were calculated for foci within each strain (Fig 2, third row; **Fig S2**). This approach allows us to extract characteristic timescales of signal fluctuations within intensity trajectories, which reflect the lifetimes of binding and dissociation events. Exponential fitting of each mean autocorrelation function gave time constants of τ = < 0.03, 0.3 and 3.3 s, reflecting short-, medium-, and long-lived binding events (**Fig S2**). For each strain and time interval after ciprofloxacin addition, the relative proportions of these binding events are plotted in Fig 2 (fourth row).

For both *umuDC*^+^ and Δ*umuDC* cells, most pol IV binding at τ positions appeared to be short-lived (Figs 2A, B). In the early stages of ciprofloxacin exposure (25–45 min) the components of the autocorrelation function were 80% short-lived (< 0.03 s, shorter than a frame of 34 ms), 10% medium (0.3 s) and 10% long-lived (3.3 s). In the later stages, (120–150 min), the proportion of medium-long lived events increased to 40%. In cells expressing UmuD(K97A) long-lived events were much more common: by the 120–150 min period medium and long-lived events comprised 80% of the autocorrelation function (Fig 2C). In stark contrast, cells expressing UmuD′ produced almost exclusively short-lived events (Fig 2D). UmuD′ appeared to supress the medium and long-lived pol IV binding events that occur in wild-type *umuDC*^+^ background following ciprofloxacin treatment.

Taken together, the results indicate that UmuD(K97A) promotes long-lived DNA binding by pol IV, whereas UmuD′ inhibits binding. The deletion of *umuDC* only marginally increases the binding lifetime of pol IV compared to *umuDC*^+^. The results demonstrate that the binding of pol IV to its substrates on the nucleoid is modulated by UmuD and UmuD′ in cells, independently of pol V formation (i.e. in cells lacking UmuC).

### Pol IV binds frequently at RecA* structures

Like UmuD and UmuD′, the RecA recombinase modulates the mutagenic activity of pol IV (Pomerantz *et al*., 2013b; Pomerantz *et al*., 2013a). *In vitro*, DNA synthesis by pol IV is error-prone when operating on D-loop substrates that mimic recombination intermediates (Pomerantz *et al*., 2013b; Pomerantz *et al*., 2013a). Pol IV is known to participate in error-prone DSB repair under a variety of circumstances (Ponder *et al*., 2005; Lovett, 2006; Williams *et al*., 2010; Shee *et al*., 2011; Rosenberg *et al*., 2012; Shee *et al*., 2012b; Shee *et al*., 2012a; Pomerantz *et al*., 2013a; Pomerantz *et al*., 2013b). *In vitro*, RecA also facilitates DNA synthesis by pol IV in replisomes (Indiani *et al*., 2013). However, it remains to be determined whether pol IV binds at RecA* *in vivo*.

We determined whether pol IV colocalises with RecA* structures by visualising the localisations of fluorescent pol IV (DinB-YPet) and a RecA* marker PAmCherry-mCI; a red fluorescent protein fusion of a monomeric C-terminal fragment of the λ repressor that retains the ability to bind RecA* in cells (Ghodke *et al*., 2019). We carried out this analysis in SSH092 cells treated with ciprofloxacin — a potent inducer of DSBs (Drlica and Zhao, 1997; Zhao *et al*., 1997) through reactive oxygen species-dependent and -independent pathways (Goswami *et al*., 2014). Live-cell photoactivatable localisation microscopy (PALM) of SSH092 cells treated with ciprofloxacin was performed by collecting images in both channels every 5 min over a period of 3 h following introduction of ciprofloxacin at time point t = 0 min. At each time point, a new field-of-view was recorded.

Following ciprofloxacin treatment, cells typically contained multiple mCI foci (Fig 3A). At later time points, some cells contained more elongated “bundle” structures as described previously (Ghodke *et al*., 2019). We next determined the percentage of DinB-YPet foci that colocalised with mCI foci and bundle-like structures (Fig 3B). Prior to the introduction of ciprofloxacin, mCI foci were rarely formed in cells during normal metabolism (< 0.1 mCI foci per cell) consistent with our previous study (Ghodke *et al*., 2019). Unsurprisingly, we did not detect colocalisation of pol IV with the RecA* probe in untreated cells. Upon introduction of ciprofloxacin to the flow chamber, colocalisation remained low during the early phase of the SOS response (i.e., between 0–45 min after treatment). From 45 min after the introduction of ciprofloxacin, pol IV exhibited extensive colocalisation (10–40%) with mCI in cells. This extensive colocalisation persisted into the late stages of SOS (up to 180 min after treatment). We have previously noted that most of the mCI foci form at locations distal to the replisome in UV-irradiated cells (Ghodke *et al*., 2019). Notably, pol IV foci also mainly form at sites distinct from replisome markers.

**Figure 3.**
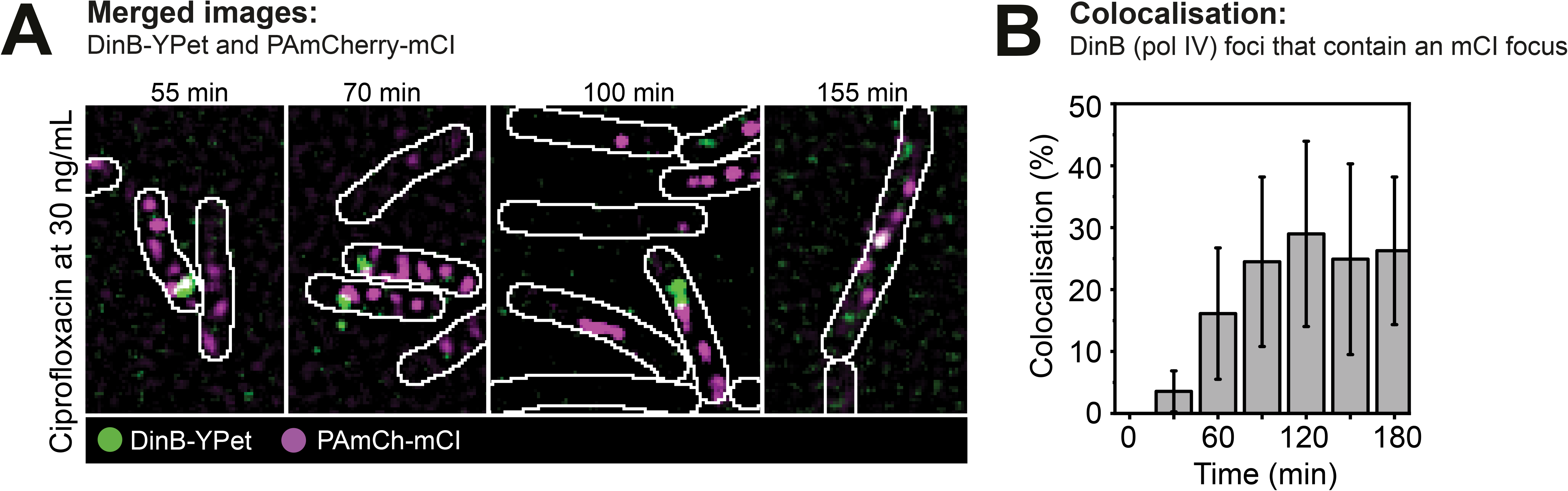
Colocalisation measurement between DinB and mCI after ciprofloxacin treatment. (A) Merged images of discoidal filtered DinB-YPet (green) and PAmCherry-mCI (magenta) at 55, 70, 100 and 155 min after ciprofloxacin addition. (B) Colocalisation percentage of DinB with mCI. Time points are binned over 30 min. Error bar represents standard deviation of biological quadruplicates.

### RecA* promotes the binding of pol IV to the nucleoid

Having observed that a large proportion of pol IV colocalises with the RecA* probe mCI, we next set out to determine whether RecA* structures could recruit pol IV to the DNA. To isolate the effects of RecA* formation from other effects introduced by exogenous DNA damage, we utilised a RecA mutant, RecA(E38K), which is able to constitutively induce SOS and high rates of pol V-dependent mutagenesis in cells (Cazaux *et al*., 1993; Wang *et al*., 1993; Ennis *et al*., 1995; Gruenig *et al*., 2008), suggestive of RecA* structures being formed in the absence of endogenous DNA damage.

Using surface plasmon resonance (SPR) as previously described (Ghodke *et al*., 2019), we observed that RecA(E38K) forms filaments on ssDNA *in vitro* (**Figs S3A, B**). Stable association of RecA(E38K) required the presence of ATPγS suggesting that RecA(E38K) forms filaments (**Fig S3B**). Additionally, RecA(E38K) filaments on ssDNA are competent to cleave LexA (**Fig S4**), suggesting that RecA(E38K) forms RecA*-like structures on ssDNA (Burckhardt *et al*., 1988; Simmons *et al*., 2008). However, in the absence of DNA damage, we expect exposed ssDNA substrates for RecA(E38K) binding to occur infrequently. Therefore, we additionally tested whether constitutive SOS signalling may occur due to constitutive RecA(E38K)-dsDNA filament formation. To that end, we tested the ability of RecA(E38K) to form filaments on a 60-mer dsDNA substrate. We found that RecA(E38K) binds readily to dsDNA (**Figs S3C, D**) and that incubation of dsDNA plasmid substrates with RecA(E38K) promoted LexA cleavage (**Fig S4**), indicating that RecA(E38K) also forms RecA*-like structures on dsDNA (Burckhardt *et al*., 1988; Simmons *et al*., 2008).

Together, these results allowed us to establish conditions where we could now probe the binding of pol IV to constitutive RecA filaments, even in the absence of exogenous DNA damage, in live cells. We carried out single-molecule imaging of *dinB-YPet dnaX-mKate2* cells carrying wild-type or mutant alleles of *lexA* (encoding the SOS-response repressor LexA) and *recA* (encoding the recombinase RecA). Three strains were examined: i) cells with wild-type *lexA* and *recA* alleles (EAW643, *dinB-YPet dnaX-mKate2 lexA^+^ recA*^+^, Table 1), ii) cells that constitutively express high levels of DinB-YPet (and all other SOS-induced proteins) even in the absence of exogenous DNA damage (Simmons *et al*., 2008); RW1594, *dinB-YPet dnaX-mKate2 lexA*[Def] *recA*^+^, Table 1) and iii) cells that both produce high levels of DinB-YPet and constitutively formed RecA*-like structures (RW1598, *dinB-YPet dnaX-mKate2 lexA*[Def] *recA*[E38K], Table 1) (Cazaux *et al*., 1993; Wang *et al*., 1993; Ennis *et al*., 1995; Gruenig *et al*., 2008). Although cells carrying the *recA*(E38K) allele are constitutive for SOS induction, our previous study of UmuC-mKate2 cells suggested that this induction only operates at ∼50% of maximum – expression of UmuC-mKate2 could be further induced by UV irradiation, whereas this was not the case for *lexA*(Def) cells (Robinson *et al*., 2015) We therefore included the additional *lexA*(Def) allele in RW1598 so that the intracellular concentration of pol IV would match that of RW1594 cells.

We set out to determine if the presence of RecA*-like structures formed by RecA(E38K) is sufficient to recruit pol IV to the nucleoid in cells. We recorded burst acquisitions of DinB-YPet motions in the three strains (300 images of 34 ms exposure time, total length of 10.2 s). For each movie, a corresponding image of the replisome marker τ-mKate2 was also captured (see **Figs S2A, B** for imaging sequence). As expected, cells with wild-type *lexA* and *recA* alleles produced few pol IV foci (Fig 4A) (Henrikus *et al*., 2018b). Cells that carried the SOS-constitutive *lexA*(Def) allele and the wild-type *recA* allele produced a relatively high level of DinB-YPet signal, but produced few foci (Fig 4B) (Henrikus *et al*., 2018b). This result is consistent with our previous study in which we concluded that binding is triggered by the presence of damage on the DNA, as opposed to mass action-driven exchange brought on by increased intracellular concentrations of pol IV (Henrikus *et al*., 2018b). In contrast to both *recA*^+^ strains, cells carrying both the *lexA*(Def) allele and the RecA*-constitutive *recA*(E38K) allele produced both high DinB-YPet signal and readily visible foci (Fig 4C). These results suggest two possibilities. First, in the absence of ciprofloxacin-induced double strand breaks, *lexA*(Def) *recA*(E38K) cells might produce some kind of DNA structures that serve as substrates for pol IV (and are not present in *lexA*(Def) cells carrying wild-type RecA). Second, nucleoid-associated RecA(E38K) assemblies might themselves acts as binding sites for pol IV in *recA*(E38K) cells.

**Figure 4.**
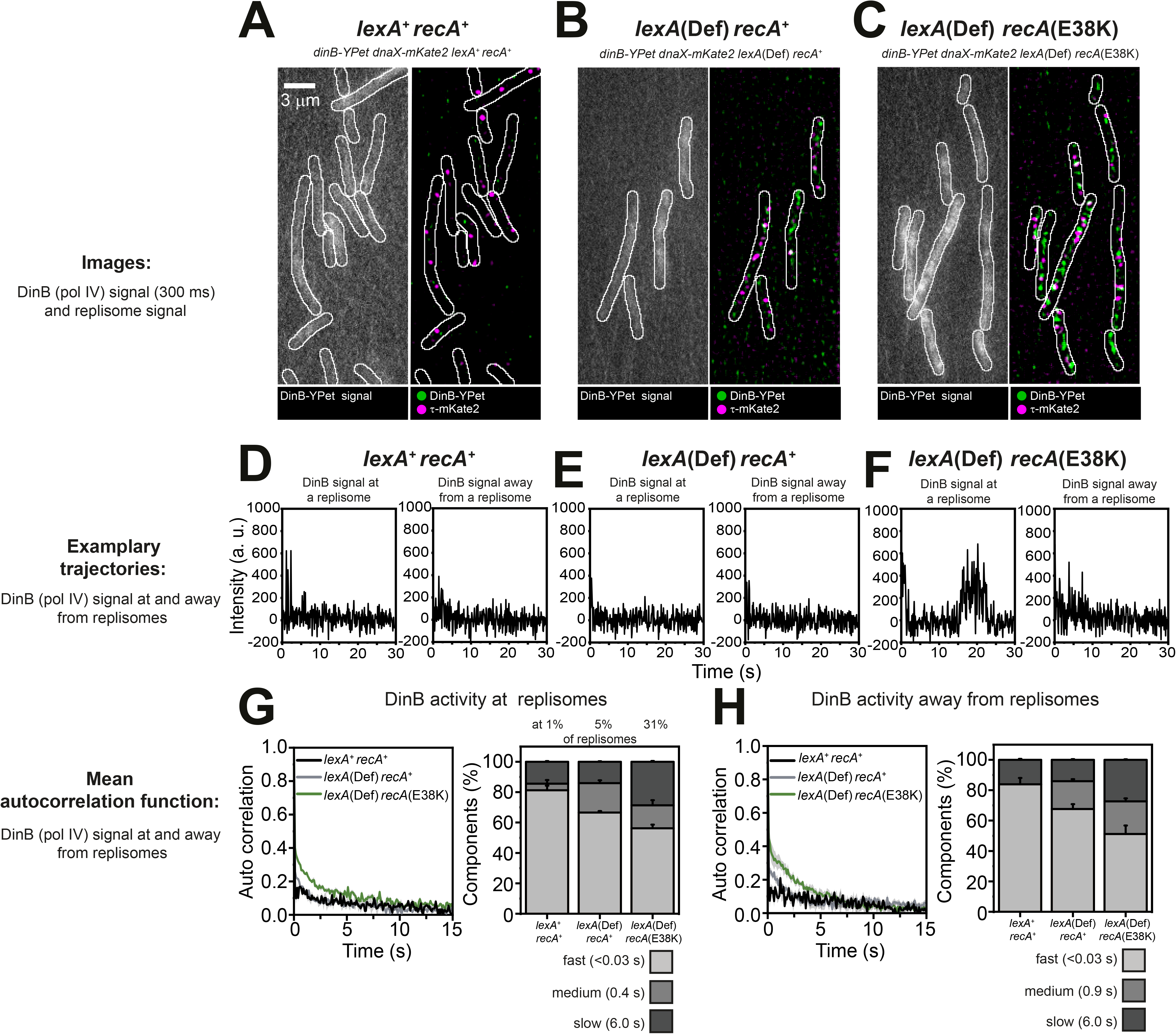
Binding activity of DinB at and away from replisomes in different *lexA* and *recA* mutants. (A) Images of DinB and DnaX signal in *lexA*^+^ *recA*^+^. Left: Projection of DinB signal consistent with 300 ms exposure times. Right: Merged images of discoidal filtered DinB (green) and DnaX signal (magenta). (B) similar to (A), however in *lexA*(Def) *recA*^+^. (C) similar to (A), however in *lexA*(Def) *recA*(E38K). (D) Left: DinB signal at a replisome in *lexA*^+^ *recA*^+^. Right: DinB signal away from replisome in *lexA*^+^ *recA*^+^. (E) Left: DinB signal at a replisome in *lexA*(Def) *recA*^+^. Right: DinB signal away from replisome in *lexA*(Def) *recA*^+^. (F) Left: DinB signal at a replisome in *lexA*(Def) *recA*(E38K). Right: DinB signal away from replisome in *lexA*(Def) *recA*(E38K). (G) Mean autocorrelation function showing DinB activity at replisomes in *lexA*^+^ *recA*^+^ (black line), *lexA*(Def) *recA*^+^ (grey line) and *lexA*(Def) *recA*(E38K) (green line). Error bars represent standard error of the mean over > 100 trajectories. (H) Mean autocorrelation function showing DinB activity away from replisomes in *lexA*^+^ *recA*^+^ (black line), *lexA*(Def) *recA*^+^ (grey line) and *lexA*(Def) *recA*(E38K) (green line). Error bars represent standard error of the mean > 100 trajectories.

We therefore directly tested whether RecA(E38K) interacts with pol IV on filaments assembled dsDNA *in vitro*. Using an identical SPR experimental setup as described above, we assembled RecA(E38K) on a 60-mer dsDNA substrate (**Figs S3C, D**). We found that pol IV associates with RecA(E38K)-ATPγS filaments formed on dsDNA (**Fig S3E**), producing a much stronger response than measurements in which pol IV was exposed to dsDNA in the absence of RecA(E38K) (**Fig S3F**). Unfortunately, despite our attempts to further optimise the assay, non-specific binding of pol IV to the chip surface hampered our attempts to extract binding parameters from the sensorgrams. Nevertheless, these results clearly demonstrate that the association of pol IV with the nucleoid is promoted by the presence of RecA*-like structures.

Returning to the live-cell single-molecule data, we next examined fluctuations in the DinB-YPet signals that occur as pol IV binds to, or dissociates from, binding sites on the nucleoid. We monitored pol IV binding events within cells, both close to and away from τ foci. Intensity trajectories for DinB-YPet in *lexA*^+^ *recA*^+^ cells and *lexA*(Def) *recA*^+^ cells predominantly showed short-lived spikes (< 1s; Figs 4D, E), indicative of transient pol IV binding events (milliseconds timescale). In contrast, trajectories for DinB-YPet in *lexA*(Def) *recA*(E38K) cells often included binding events that were much longer lived (1–10 s, Fig 4F), indicative of pol IV binding to its target for longer periods (seconds timescale).

To comprehensively assess pol IV binding lifetimes across all intensity trajectories, mean autocorrelation functions were calculated for each set of trajectories (**Figs S2D–F**). Fitting of each autocorrelation function give time constants τ = < 0.03, 0.4 and 6.0 s, reflecting short-, medium-, and long-lived binding events (Figs 4G, H; **S2G, H**). For *lexA*^+^ *recA*^+^ cells in the absence of ciprofloxacin, only 1% of τ positions showed evidence of pol IV binding events (Fig 4G, right panel). The normalised mean autocorrelation function for *lexA*^+^ *recA*^+^ cells was of low amplitude (0.16 at Δ*t* = 1 frame, Fig 4G, black line), indicative of there being relatively few long-lived binding events at replisomes across the different trajectories (Henrikus *et al*., 2018b). The *lexA*(Def) *recA*^+^ background marginally increased pol IV binding activity with 5% of τ foci showing by DinB-YPet binding (Fig 4G, right panel) (Henrikus *et al*., 2018b). The autocorrelation function remained of low amplitude (0.3 at Δ*t* = 1 frame, Fig 4G, grey line), indicating that few long-lived pol IV binding events occurred at τ positions in the *lexA*(Def) *recA*^+^ background. In contrast, *lexA*(Def) *recA*(E38K) cells exhibited a strong increase in pol IV binding activity, both close to and away from τ foci; 31% of τ positions had a pol IV binding event (Fig 4G, right panel). The amplitude of the autocorrelation function was also increased (0.4 at Δ*t* = 1 frame, Fig 4G, green line), indicating that long-lived binding events occurred close to replisome markers much more frequently. The decay rate of the autocorrelation function had two longer timescale components (Fig 4G, right panel: τ_m_ = 0.4 s with an amplitude of 15% and τ_l_ = 6.0 s with an amplitude of 29%), suggesting that pol IV typically binds near τ foci for periods of a few seconds in the *recA*(E38K) background. When analysing the binding behaviour of pol IV away from τ positions in these three backgrounds, similar results were obtained (Fig 4H).

## Discussion

In this study, we arrived at four conclusions: i) UmuD promotes the binding of pol IV to the nucleoid, at replisomal and non-replisomal sites; ii) UmuD′ inhibits the binding of pol IV to the nucleoid, at both replisomal and non-replisomal sites; iii) pol IV frequently colocalises with the RecA* probe mCI; iv) RecA*-like structures strongly promote the binding of pol IV to the DNA. These results lead us to infer that RecA*-like structures can recruit pol IV to the nucleoid. Following ciprofloxacin treatment, this pol IV-RecA* interaction might recruit pol IV to carry out repair synthesis at DSB repair intermediates. Furthermore, the RecA* mediated cleavage of UmuD, a biochemical switch that has long been known to regulate pol V activation, also regulates the binding of pol IV to the nucleoid. The results provide direct evidence for both RecA and UmuD acting as regulatory factors for pol IV *in vivo*, as proposed previously (Godoy *et al*., 2007; Indiani *et al*., 2013; Pomerantz *et al*., 2013a; Pomerantz *et al*., 2013b; Cafarelli *et al*., 2014).

### UmuD_2_ and UmuD′_2_ as regulators of pol IV

A previous study suggested that both UmuD_2_ and UmuD′_2_ bind to pol IV and modulate its mutagenic activity (Godoy *et al*., 2007). Moreover, *in vitro* experiments have suggested that full-length UmuD binds to the replicative polymerase, α, and destabilises its interaction with the sliding clamp, β, thus facilitating other polymerases, such as pol IV, to access the replisome (Sutton *et al*., 1999; Silva *et al*., 2012).

Here we show that UmuD(K97A) increases the number of pol IV foci and increases the binding time of pol IV at the nucleoid. In contrast, UmuD′ inhibits nucleoid binding by pol IV. During the first stage of the SOS response, most UmuD is present as full-length UmuD_2_ (Robinson *et al*., 2015). This would promote long-lived binding of pol IV to DNA and support high-fidelity DNA synthesis. Based on rates of pol IV-dependent DNA synthesis measured *in vitro* (3–5 nt s^−1^; (Wagner *et al*., 2000)), binding events lasting a few seconds, such as those observed during this study, could permit the incorporation of tens of nucleotides. In cells lacking *umuDC*, the operon encoding for pol V, we observed increased colocalisation between pol IV and the replisome marker, however nucleoid-binding was shorter-lived than in cells expressing UmuD(K97A). These effects of UmuD and UmuD′ were observed in strains lacking UmuC, indicating that the changes in replisome colocalization and nucleoid-binding lifetime did not arise from differences in competition for binding sites with pol V.

This work allows us to propose the following model for pol IV activity in the context of the SOS response. Cells experiencing extensive DNA damage trigger the full extent of the SOS response, leading to the formation of UmuD′ at late time points after DNA damage. At this point, the cell enters a mutagenic phase. The highly error-prone polymerase pol V Mut becomes active and pol IV, now in the absence of UmuD, introduces –1 frameshift mutations. At the same time pol IV binding becomes infrequent and short-lived in the presence of UmuD′, consistent with an earlier observation that UmuD′ reduces the frequency of adaptive mutagenesis (Godoy *et al*., 2007). Thus, while pol IV is error-prone in the presence of UmuD′, mutagenesis would be kept in check by pol IV having reduced access to substrates. This mechanism of UmuD cleavage restricting mutagenesis is in line with the multiple mechanisms that have evolved to restrict the mutagenic activity of pol V (Jaszczur *et al*., 2016). Interestingly, colocalization between pol IV and τ is highest in cells that lack UmuD and UmuC altogether (Δ*umuDC*). One possibility is that in wild-type *umuDC*^+^ cells pol V competes with pol IV for binding to replisome-proximal binding sites, however this explanation seems unlikely for two reasons: 1. pol IV-τ colocalization is low in cells that express UmuD′, but lack UmuC and therefore cannot produce pol V (Fig 1D); 2. fluorescently labelled pol V colocalises with replisomes even less frequently than pol IV does (Robinson *et al*., 2015). Another explanation, which is more consistent with the data, is that the accumulation of UmuD′ in response to treatment with DNA damaging agents inhibits the binding of pol IV at replisome-proximal sites in wild-type (*umuDC*^+^) cells.

### Pol IV binds to RecA* structures

The high degree of colocalisation we observed between pol IV and the RecA* probe when treating with ciprofloxacin, together with the binding of pol IV to RecA*-like structures *in vitro* and *in vivo*, adds to a growing body of evidence supporting the participation of pol IV in homologous recombination (Ponder *et al*., 2005; Lovett, 2006; Williams *et al*., 2010; Shee *et al*., 2011; Rosenberg *et al*., 2012; Shee *et al*., 2012b; Shee *et al*., 2012a; Pomerantz *et al*., 2013a; Pomerantz *et al*., 2013b; Moore *et al*., 2017). In ciprofloxacin-treated cells, pol IV colocalises with the RecA* probe (this study) far more frequently than it colocalises with the replisome marker τ (Henrikus *et al*., 2018b). Ciprofloxacin is a DNA gyrase inhibitor, which generates DSBs (Zhao *et al*., 1997) and rapidly halts DNA synthesis (Deitz *et al*., 1966; Snyder and Drlica, 1979). Defects in DSB processing strongly suppress both pol IV up-regulation and focus formation (Henrikus *et al*., 2019b). Interestingly, *in vitro*, pol IV is capable of associating with RecA(E38K)-ATPγS filaments formed on dsDNA. These filaments are competent of LexA cleavage, indicative of RecA*-like activity. *In vivo* in the absence of DNA damage, pol IV forms foci in a *recA*(E38K) mutant background, suggestive of pol IV binding to RecA(E38K) filaments, which presumably form predominantly on dsDNA. In wild-type cells, an interaction between pol IV and RecA* may well facilitate the recruitment of pol IV to homologous recombination intermediates, or indeed any substrates where amenable RecA* structures form.

The results presented here indicate that in ciprofloxacin-treated cells pol IV primarily forms foci at sites of RecA* structures. Together with the observation that inhibition of DSB resection almost completely eliminates pol IV focus formation in ciprofloxacin-treated cells (Henrikus *et al*., 2019b), this suggests that pol IV predominantly acts at double-strand break repair intermediates in ciprofloxacin-treated cells, where its most likely role is the extension of D-loops during repair synthesis (Indiani *et al*., 2013; Pomerantz *et al*., 2013b; Pomerantz *et al*., 2013a). The association of pol IV with RecA has also been observed outside the context of RecA* structures (Godoy *et al*., 2007) and is proposed to stimulate pol IV-dependent TLS *in vitro* (Cafarelli *et al*., 2014). Further research is required to determine whether the pol IV-RecA* interaction plays a role in modulating pol IV activities within pathways other than double-strand break repair.

### Experimental procedures

#### Strain construction, plasmid construction and transformations

SSH007 is a two-colour strain (*dinB-YPet dnaX-mKate2* Δ*umuDC*) derived from EAW643 (*dinB-YPet dnaX-mKate2*). It was made by replacing the wild-type *umuDC^+^* gene of EAW643 with Δ*umuDC*::Cm^R^ from RW880 *via* P1 transduction. Colonies were selected by testing for chloramphenicol resistance.

To investigate the influence of UmuD mutants on pol IV activity, SSH007 was complemented with plasmids that express UmuD(K97A) (pJM1243) or UmuD’ (pRW66).

SSH092 was made by transformation; EAW633 (*dinB-YPet*) carries the pJMuvrA-PAmCherry-mCI vector (see **Supplementary Notes** for sequence). The PAmCherry-mCI gene block was commercially synthesised and the sequence was verified (IDT gene block). The gene block was introduced into pSC101 (Churchward *et al*., 1984) using the *Apa*I and *Sac*II restriction sites.

RW1598 was made by P1 transduction of *recA730 srlD300*::Tn*10* from RW244 into RW1594, selecting for TetR. Colonies were then screened for constitutive UmuD cleavage using Western blotting. *recA* and *srlD* are about 90% linked.

pJM1243 was made by chemically synthesizing an *E.coli* codon optimised *umuD*(K97A) gene that was cloned into the low-copy spectinomycin resistant vector, pSC101 (Churchward *et al*., 1984), as *Hin*dIII-*Eco*RI fragment. UmuD(K97A) expression is LexA-regulated. Similarly, pRW66 was made by introducing the *umuD*’ gene into pSC101 (Churchward *et al*., 1984).

#### Fluorescence microscopy and imaging protocols

For all experiments except for experiments including imaging of PAmCherry-mCI, wide-field fluorescence imaging was performed on an inverted microscope (IX-81, Olympus with a 1.49 NA 100× objective) in an epifluorescence configuration, as described previously (Robinson *et al*., 2015). Continuous excitation is provided using semidiode lasers (Sapphire LP, Coherent) of the wavelength 514 nm (150 mW max. output) and 568 nm (200 mW max. output). τ-mKate2 was imaged using yellow excitation light (λ = 568 nm) at high intensity (2750 W cm^−2^), collecting emitted light between 610–680 nm (ET 645/75m filter, Chroma) on a 512 × 512 pixel^2^ EM-CCD camera (C9100-13, Hamamatsu). For DinB-YPet time-lapse imaging, we used green excitation (λ = 514 nm) at lower power (240 W cm^−2^), collecting light emitted between 525–555 nm (ET540/30m filter, Chroma).

For experiments including imaging of PAmCherry-mCI, imaging was conducted on an inverted microscope (Nikon Eclipse-Ti), equipped with a 1.49 NA 100× objective and a 512 × 512 pixel^2^ Photometrics Evolve CCD camera (Photometrics, Arizona, US). NIS-Elements equipped with JOBS module was used to operate the microscope (Nikon, Japan). Continuous excitation is provided using semidiode lasers of the wavelength 405 nm (OBIS, Coherent, 200 mW max. output), 514 nm (Sapphire LP, Coherent, 150 mW max. output) and 568 nm (Sapphire LP, Coherent, 200 mW max. output). DinB-YPet was imaged using green excitation (λ = 514 nm) at lower power (∼2200 W cm^−2^), collecting light emitted between 535–550 nm (ET535/30m filter, Chroma). PAmCherry-mCI was imaged by simultaneous illumination with the activation laser 405 nm (1–5 W cm^−2^) and 568 nm readout laser (540 W cm^−2^), a PALM (photoactivation localisation microscopy) acquisition protocol, collecting emitted light from 590 nm (ET590LP, Chroma).

Burst acquisitions (movies of 300 × 34 ms frames, continuous excitation with 514 nm light; each frame at 80 W cm^−2^) were collected to characterise DinB-YPet binding kinetics; followed by a set of two images (bright-field [34 ms exposure]; mKate2 fluorescence [100 ms exposure]). Data were recorded from 20–45 min, from 55–85 min and from 120–180 min after ciprofloxacin treatment (30 ng mL^−1^). Time-lapse movies were recorded to visualise changes in DinB-YPet binding activity and measure colocalisation with the replisome marker. Sets of three images were recorded (bright-field [34 ms exposure], YPet fluorescence [50 ms exposure]; mKate2 fluorescence [100 ms exposure]) at an interval of 10 min for 3 h. All images were analysed with ImageJ (Schneider *et al*., 2012).

Time-sampling of DinB-YPet and PAmCherry-mCI expressing cells were performed as follows: First, the bright-field image was taken with 100 ms exposure time. Then, a PALM acquisition protocol (simultaneous illumination with the activation laser 405 [1–5 W cm^−2^] and 568 nm readout laser [540 W cm^−2^] for 200 frames taken every 100 ms) was used to image PAmCherry-mCI. Third, DinB-YPet was detected using 512 nm laser (50 ms exposure time at ∼2200 W cm^−2^). The experiment was performed over 3 h, time points were sampled at an interval of 5 min. At each time point, a new field-of-view was sampled to minimise laser-induced damage.

To image DinB-YPet and PAmCherry-mCI, sets of three acquisitions were recorded (bright-field [100 ms exposure], YPet fluorescence [50 ms exposure]; PAmCherry fluorescence [simultaneous illumination with the activation laser 405 and 568 nm readout laser for 200 frames each with 100 ms exposure]). This protocol was only executed once for a field-of-view to minimise laser damage. Consequently, each time point shows a new set of cells. The experiment was conducted over 3 h, an image was taken every 5 min.

#### Flow cell design

All imaging was carried out on cultures growing in home-built flow cells. Imaging was carried out in quartz-based flow cells, similar to those used in our previous study (Henrikus *et al*., 2018b). These flow cells were assembled from a no. 1.5 coverslip (Marienfeld, reference number 0102222 or 0107222), a quartz top piece (45 × 20 × 1 mm^3^) and PE-60 tubing (Instech Laboratories, Inc.). Prior to flow cell assembly, coverslips were silanized with aminopropyltriethoxy silane (APTES; Alfa Aeser). First, coverslips were sonicated for 30 min in a 5 M KOH solution to clean and activate the surface. The cleaned coverslips were rinsed thoroughly with MilliQ water, then treated with a 5% (v/v) solution of APTES in MilliQ water. The coverslips were subsequently rinsed with ethanol and sonicated in ethanol for 20 s. Afterwards, the coverslips were rinsed with MilliQ water and dried in a jet of N_2_. Silanised slides were stored under vacuum prior to use.

To assemble each flow cell, polyethylene tubing (BTPE-60, Instech Laboratories, Inc.) was glued (BONDiT B-482, Reltek LLC) into two holes that were drilled into a quartz piece. After the glue solidified overnight, double-sided adhesive tape was stuck on two opposite sides of the quartz piece to create a channel. Then, the quartz piece was stuck to an APTES-treated coverslip. The edges were sealed with epoxy glue (5 Minute Epoxy, DEVCON home and Epoxy Adhesive, 5 Minute Everyday, PARFIX). Each flow cell was stored in a desiccator under mild vacuum while the glue dried. Typical channel dimensions were 45 × 5 × 0.1 mm.

#### Setup of flow cell experiments

For all imaging experiments, cells were grown at 37 °C in EZ rich defined medium (Teknova) that contained 0.2% (w/v) glucose. EAW643, RW1594 and RW1598 cells were grown in the presence of kanamycin (25 μg mL^−1^), SH001 cells were grown in the presence of chloramphenicol (25 μg mL^−1^), SSH007 cells carrying pJM1243 or pRW66 were grown in the presence of spectinomycin (50 μg mL^−1^). Cells carrying PAmCherry-mCI were also grown in the presence of spectinomycin (50 μg mL^−1^). Cells were loaded into flow cells, allowed a few minutes to associate with the APTES surface, then, loosely associated cells were removed by pulling through fresh medium. The experiment was then initiated by switching the medium to a medium that contains 30 ng mL^−1^ ciprofloxacin (for cells carrying plasmids: 50 μg mL^−1^ spectinomycin was added). A flow rate of 50 μL min^−1^ was applied during the experiment to allow a constant nutrient and oxygen supply by using a syringe pump.

#### Proteins

The wild-type *E. coli* RecA protein was purified as described (Craig and Roberts, 1981). The RecA concentration was determined using the extinction coefficient ε_280_ = 2.23 × 10^4^ M^-1^ cm^-1^ (Craig and Roberts, 1981).

The *E. coli* RecA(E38K) protein was purified as previously described (Cox *et al*., 2003) with the following modifications. After washing the protein pellet with R buffer plus 2.1 M ammonium sulfate, the pellet was resuspended in R buffer plus 1 M ammonium sulfate. The sample was loaded onto a butyl-Sepharose column and washed with 1.5 column volumes of R buffer plus 1 M ammonium sulfate. It was then eluted with a linear gradient from R buffer plus 1 M ammonium sulfate to R buffer, carried out over 5 column volumes. Peak fractions were identified by SDS-PAGE analysis and pooled. The protein was loaded onto a hydroxyapatite column as done previously, but with the linear gradient from 10–500 mM P buffer. The fractions were dialyzed against R buffer plus 50 mM KCL and 1 mM dithiothreitol three times. The fractions were loaded onto a Source 15S column and washed with R buffer plus 50 mM KCl and 1 mM dithiothreitol until the UV trace receded from peak. Next, the pool was loaded onto a Source 15Q column and eluted with a linear gradient from 0.05–1 M KCl over 25 column volumes. Peak fractions were identified as above and pooled. A DEAE-Sepharose column was not used. Protein in this pool was precipitated by the addition of equal volume of 90% saturated ammonium sulfate. The precipitate was stirred and then spun down at 13,000 rpm for 30 min. The pellet was resuspended in R buffer plus 1 M ammonium sulfate, stirred for an hour, and then spun down again. This protein was loaded onto a butyl-Sepharose column and eluted in a gradient from R buffer and 1 M ammonium sulfate to R buffer. The fractions were identified, pooled, and concentrated using GE Vivispin 20 10K MWCO centrifuge filter concentrating units. The protein was flash frozen in liquid nitrogen and stored at –80 °C. The concentration was determined as above. No exonuclease or other endonuclease activities were detected.

Pol IV (*dinB*) coding sequence was cloned into NcoI and BamHI sites of pET16b to generate a native pol IV expression construct. *E. coli* strain Turner/pLysS (Novagen) carrying the expression construct was grown in LB medium supplemented with 20 μg/ml chloramphenicol and 100 μg ml^−1^ ampicillin. Expression of pol IV was induced by adding IPTG to 1 mM and growing for 3-4 h at 30°C. Collected cells (∼20 g) were resuspended in 50 mL of lysis buffer (50 mM Tris-HCl, pH 7.5, 1 M NaCl, 10% sucrose, 2 mM DITHIOTHREITOL, 1 mM EDTA and protease inhibitor cocktail). Cells were lysed by lysozyme (2 mg/mL) and the clarified extract was collected following centrifugation at 15000 x g for 30 min. Pol IV was then precipitated by ammonium sulfate added to 30% saturation and stirring for 10 min. The precipitate was subjected to gel-filtration in GF-buffer (20 mM Tris-HCl, pH 7.5, 1 M NaCl, 0.1 mM EDTA, 1 mM DITHIOTHREITOL) using a GE Healthcare Superdex-75 XK-26/60 gel filtration column. Pol IV fractions were pooled, dialyzed overnight in PC-buffer (20 mM Tris-HCl, pH 7.5, 0.1 mM EDTA 1 mM DITHIOTHREITOL, 10% glycerol), containing 200 mM NaCl and then subjected to phosphocellulose chromatography (P-11, Whatman). After washing extensively with PC-buffer + 200 mM NaCl, pol IV was eluted with a linear gradient of 200–500 mM NaCl. Fractions containing native pol IV (> 99% pure) were pooled and stored at –70 °C.

#### Surface Plasmon Resonance (SPR) experiments

SPR experiments were conducted on BIAcore T200 istrument (GE Healthcare) using streptavidin (SA) coated sensor chips, probing the formation of RecA structures (assembled from RecA[E38K]) on ssDNA and dsDNA. Experiments were carried out at 20 °C at a flow rate of 5 μL min^−1^. As described previously (Ghodke *et al*., 2019), SA chips were activated and stabilised, single-stranded biotinylated 71-mer poly-dT oligonucleotide bio-(dT)_71_ was immobilised, followed by RecA(E38K) filament assembly (**Figs S3A, B**). RecA(E38K) filaments were assembled on bio-(dT)_71_ by injecting 1 μM RecA(E38K) in SPR^RecA(E38K)^ buffer (20mM Tris-HCl, pH 8.0, 10 mM KCl, 10 mM MgCl_2_, 0.005% surfactant P20 and 0.5 mM dithiothreitol) supplemented with 1 mM adenosine 5’-(γ-thio) triphosphate (ATPγS) at 10 μL min^−1^ for 400 s. Similarly, biotinylated dsDNA was immobilised (as previously described (Ghodke *et al*., 2019)), followed by RecA(E38K) filament assembly (**Figs S3C, D**). RecA(E38K) filaments were assembled on dsDNA (sequence: 5’-TCC TTT CGT CTT CAA AGT TCT AGA CTC GAG GAA TTC TAA AGA TCT TTG ACA GCT AGC CAG-3’, 5’ end is biotinylated) by injecting 1 μM RecA(E38K) in SPR^RecA(E38K)^ buffer (20mM Tris-HCl, pH 8.0, 10 mM KCl, 10 mM MgCl_2_, 0.005% surfactant P20 and 0.5 mM dithiothreitol) supplemented with 0.5 mM ATPγS at 5 μL min^−1^ for 500 s. Then, SPR^RecA(E38K)^ supplemented with 0.5 or 1 mM ATPγS buffer was flowed in at 5 μL min^−1^ for 2,500 s, in order to stabilise the formed filaments. From 3,000 s, 1 μM RecA(E38K) in SPR^RecA(E38K)^ buffer supplemented with 0.5 mM ATPγS was injected at a flow rate of 5 μL min^−1^ for 4,200 s.

Pol IV association with RecA(E38K)-dsDNA filaments was observed by injecting 0.65 μM pol IV in SPR^RecA(E38K)^ buffer supplemented with 0.5 mM ATPγS for 220 s at 5 μL min^−1^, monitoring pol IV association (**Fig S3E**). From 220 s, buffer containing 0.5 mM ATPγS was flowed in at 5 μL min^−1^ and fast dissociation of pol IV was observed. Similarly, pol IV association with dsDNA was monitored, giving a lower response curve (**Fig S3F**). We also observed non-specific binding of pol IV to the chip surface, making it impossible to measure binding kinetics of pol IV.

The surface was regenerated as previously reported (Ghodke *et al*., 2019). Furthermore, the SPR signal were corrected using a flow cell without immobilised bio-(dT)_71_ or dsDNA and corrected for the amount of immobilised RecA(E38K) (Ghodke *et al*., 2019). Ghodke *et al*. utilised this assay to monitor the binding kinetics of mCI at RecA-ssDNA filament (Ghodke *et al*., 2019).

#### DNA substrates for ATPase and LexA cleavage assay

M13mp18 cssDNA was purified as previously described (Neuendorf and Cox, 1986), and M13mp18 cdsDNA was prepared as previously described (Messing, 1983; Neuendorf and Cox, 1986; Haruta *et al*., 2003). The M13mp18 nicked dsDNA (from here onward called pEAW951) was prepared by nicking with DNaseI according to manufacturer’s recommendations. All DNA concentrations are given in terms of total nucleotides.

#### ATPase assay

ATP hydrolysis of wild-type RecA and RecA(E38K) on nicked cdsDNA was measured using a spectrophotometric enzyme assay (Lindsleys and Cox, 1990; Morrical and Cox, 1990). ATP regeneration from phosphoenolpyruvate and ADP was coupled to the oxidation of NADH, which was monitored by the decrease in absorbance of NADH at 380 nm. 380-nm light was used so that the signal remained within the linear range of the spectrophotometer during the experiment. The assays were carried out on a Varian Cary 300 dual beam spectrophotometer equipped with a temperature controller and a 12-position cell changer. The cell path length and band pass were 0.5 cm and 2 nm, respectively. The NADH extinction coefficient at 380 nm of 1.21 mM^−1^ cm^−1^ was used to calculate the rate of ATP hydrolysis.

The reactions were carried out at 37 °C in a buffer containing 25mM Tris-Ac (80% cation, pH 7.5), 3 mM potassium glutamate, 10 mM magnesium acetate, 5% (w/v) glycerol, 1mM dithiothreitol, an ATP regeneration system (10 units ml^−1^ pyruvate kinase, 3 mM phosphoenolpyruvate), and a coupling system (2 mM NADH and 10 units ml^−1^ lactate dehydrogenase). The concentration of DNA (pEAW951 nicked cdsDNA) was 5 µM. One cuvette was a blank control that contained everything except the DNA (volume compensated with TE). The nicked cdsDNA, buffer, and ATP regeneration system were preincubated at 37 °C for 10 min before addition of 3 mM ATP and 3 µM wild-type RecA or RecA(E38K). Data collection was then begun.

#### LexA cleavage assay

The cleavage of LexA was performed essentially as previously described (Burckhardt *et al*., 1988). Reaction mixtures (125µl) contained 40 mM Tris-HCl, pH 8.0, 10 mM MgCl_2_, 30 mM NaCl, 2 mM dithiothreitol, 3 µM of M13mp18 circular single-stranded DNA or pEAW951 nicked circular double-stranded DNA, 3 mM ATPγS, LexA, and RecA as noted. Reactions were incubated at 37 °C for 10 min before addition of LexA. The reaction products were separated and visualized by 15% SDS-PAGE stained with Coomassie blue.

#### Analysis of colocalisation events of pol IV with replisomes

Foci were classed as colocalised if their centroid positions (determined using our peak fitter tool) fell within 2.18 pixels (218 nm) of each other (Henrikus *et al*., 2019a). For colocalisation analysis, we binned the data in 30 min intervals for a larger sample size per time point due to low numbers of pol IV foci per cell at exposures of 300 ms. We determined that for DinB-YPet–τ-mKate2 localisation the background of pol IV foci expected to colocalise with replisomes purely by chance is ∼4%. This was calculated by taking the area of each cell occupied by replisome foci (including the colocalisation search radius) and dividing by the total area of the cell. The value of 4% corresponds to the mean of measurements made over > 300 cells. As the number of pol IV foci changes in time, the proportion of replisome foci expected to colocalise with pol IV foci by chance also changes in time. At an exposure time of 50 ms, there are almost zero pol IV foci at the beginning of the measurement, thus there is close to zero probability that a replisome focus will colocalise with a pol IV focus by chance. At t = 30 min, chance colocalisation is expected to be 5% and at t = 120 min, the chance colocalisation is expected to be 3%. At an exposure time of 300 ms, the number of pol IV foci per cell never exceeds ∼0.6 foci per cell, thus the level of colocalisation expected to occur by chance is close to zero.

#### Analysis of pol IV binding kinetics

Replisome localisations were determined by identifying and fitting peaks from τ-mKate2 images. From the corresponding burst acquisition movie, the DinB-YPet signal at replisomes was plotted against time (trajectories of DinB-YPet activity at replisomes) from 20–45 min, from 55–85 min and from 120–180 min after ciprofloxacin treatment (**Fig S2C**). These were divided into trajectories that give and not give pol IV binding events (**Figs S2D, E**). From this, the percentage of replisomes (τ-mKate2 foci) that are visited by DinB-YPet molecules (Fig 4G, right panel) is calculated.

Only trajectories that have pol IV binding events were then used to separate pol IV binding kinetics. The autocorrelation function was applied to each of these trajectories giving signal similarities as a function of the lag time, a method that identifies time-dependent fluctuations in signal dependent on binding and dissociation of molecules. When applying the autocorrelation function to a DinB-YPet trajectory, the correlation of this trajectory with its time delayed copy is generated for various lag times. With zero lag time, the normalised correlation of a trajectory with itself is 1. The correlation of a trajectory with its time delayed copy, however, gives autocorrelation values that range from 0–1 depending on signal fluctuations. DinB-YPet molecules that are statically bound show no fluctuations in the DinB-YPet fluorescence signal over time, consistent with the signal being correlated in time. Consequently, the autocorrelation is between 0–1 for lag times after zero. In contrast, DinB-YPet molecules that are transiently associated show many fluctuations in the DinB-YPet fluorescence signal over time. The signal is not correlated in time and results in zero autocorrelation for lag times after zero.

For each time window (20–45 min, 55–85 min and 120–180 min), the mean autocorrelation function output was calculated to determine the average of DinB-YPet binding kinetics. The fast decay at t = 0 s corresponds to short components. From t > 0 s, the curve was fitted with a two-exponential function where medium or short components were identified (y=y_0_+A_1_·e^-x·τ1^+A_2_·e^-x·τ2^). Using the *in vitro* experimentally determined rate of nucleotide incorporation of pol IV as a guide (3–5 nt s^−1^ (Wagner *et al*., 2000)), the short, medium and long components are translated to no binding/short-lived binding (unproductive binding), binding events that are sufficient for the insertion of 1–2 nucleotides or ∼8.5 nucleotides, respectively.

Pol IV binding activity away from replisomes was determined as described above. Pol IV trajectories were, however sampled, from average projections of pol IV burst acquisitions in time (average projection over 100 frames, exposure time for each frame was 34 ms; total exposure 3.4 s; see Fig 2, upper row).

#### Analysis of colocalisation events of pol IV with mCI

To measure colocalisation between pol IV and mCI, we first created a maximum projection of each PAmCherry-mCI movie. Similar to the colocalisation analysis performed for pol IV with replisomes, foci were classed as colocalised if their centroid positions fell within 218 nm of each other. Chance colocalisation of pol IV with mCI is close to zero at 0 min. Chance colocalisation is increased from 50 min with ∼4%. At 100 min, the chance colocalisation is ∼15%.

## Acknowledgements

We thank John P. McDonald for the pJM1243 plasmid and Roger Woodgate for RW880, RW1594 and RW1598 strains, and pRW66 plasmid. We also thank Richard Spinks for helpful discussions. MMC was supported by grant GM32335 from the National Institute of General Medical Sciences USA. AvO was supported by a Laureate Fellowship FL140100027 from the Australian Research Council. MFG was supported by grant R35ES028343 from the National Institute of Environmental Health Sciences USA. AR was supported by Project Grant APP1165135 from the National Health and Medical Research Council and funds from the Faculty of Science, Medicine and Health and the Illawarra Health and Medical Research Institute.

## Author contributions

SSH, AEM, SJ, MMC, AMvO, HG and AR made contributions to the design of the study. SJ, MLR, PTP, EAW, HG and AR contributed resources to undertake the study. SSH, AEM, SJ, MLR, MMC, HG and AR were involved the interpretation of the data. SSH, AEM, SJ and MLR were involved in the acquisition and analysis of the data. SSH, AEM, MFG, MMC, AMvO, HG and AR wrote the manuscript.

## Figure legends

**Supplementary Figure S1. Colocalisation analysis using 50 ms exposures for DinB-YPet and number of DinB-YPet and τ-mKate2 foci per cell.** (A) Upper row: colocalisation of DinB with DnaX. Left plot compares *umuDC*^+^ (black line) with Δ*umuDC* (green line). Middle plot compares *umuDC*^+^ (black line) with Δ*umuDC* + UmuD(K97A) expressed from plasmid (green line). Right plot compares *umuDC*^+^ (black line) with Δ*umuDC* + UmuD’ expressed from plasmid (green line). Bottom row: colocalisation of DnaX with DinB. Left plot compares *umuDC*^+^ (black line) with Δ*umuDC* (green line). Middle plot compares *umuDC*^+^ (black line) with Δ*umuDC* + UmuD(K97A) expressed from plasmid (green line). Right plot compares *umuDC*^+^ (black line) with Δ*umuDC* + UmuD’ expressed from plasmid (green line). Error bars represent standard error of the mean between at least biological triplicates. (B) Number of DinB (upper plot) and DnaX foci per cell (bottom plot) in *umuDC*^+^ (black line), Δ*umuDC* (red line), Δ*umuDC* + UmuD(K97A) (yellow line) and Δ*umuDC* + UmuD’ (blue line) after ciprofloxacin treatment. Error bars represent standard error of the mean for *n* > 100 cells.

**Supplementary Figure S2. Burst acquisitions and analysis.** (A) Experimental setup. Cells are loaded in a flow cell and immobilised on a positively charged APTES glass surface. Cells were imaged before addition of ciprofloxacin and 25–45, 55–85 and 120–150 min after addition. (B) Burst acquisition sequence. Movies of DinB-YPet were recorded. The movies contain 300 frames at an exposure of 50 ms taken every 100 ms. Subsequently, an image of the τ-mKate2 channel is taken at an exposure time of 100 ms. (C) Exemplary intensity trajectories showing DinB-YPet binding at replisomes. (D) Histogram of DinB-YPet intensities at replisomes. From cut-off to 0: replisomes with no DinB-YPet binding. From cut-off to higher intensities: replisomes with DinB-YPet binding. (E) Grouping of trajectories. Trajectories that show no DinB-YPet binding at replisomes are excluded from the analysis. Trajectories that show DinB-YPet binding at replisomes are used for the analysis. (F), The mean autocorrelation function is obtained from single autocorrelation function. Each autocorrelation function stems from single intensity trajectories of a DinB-YPet binding event at replisomes. (G), Determining components of autocorrelation functions. The mean autocorrelation function is plotted in black. The autocorrelation function has short-lived components which consist of noise, short-lived and transient binding events (light grey line). Slower components retrieved from longer-lived events are fitted with a two-exponential fit (green line) which consist of medium and slow decorrelation events consistent with binding events. (H), Components of the autocorrelation function are plotted in a bar graph. Long, medium and short components are indicated by different colours: long (dark green), medium (light green), short (light grey). The error bars for long and medium components were extracted from the fit error using the two-exponential fit. The error bar from the short-lived components is equivalent to the standard error of the mean from the mean autocorrelation function at lag time 0s.

**Supplementary Figure S3. Sensorgram showing RecA(E38K) filament assembly on ssDNA and dsDNA in order to probe interactions with pol IV.** (A) Sensorgram showing the immobilisation of ssDNA, (dT)_71_, on the SA chip surface (association: dark grey phase; immobilised ssDNA: light grey phase). (B) Following ssDNA immobilisation, buffer containing 1 μM RecA(E38K) (+ 1 mM ATPγS) was flowed into the flow cell, at t = 0 min for 400 s. During this period, RecA(E38K) associated with ssDNA (blue phase), forming a RecA(E38K) filament. At 400 s, buffer containing 1 mM ATPγS was flowed into the flow cell. RecA(E38K) dissociates from the surface (yellow phase). From 1,400 s, RU units are constant, consistent with stabilised RecA(E38K) filaments. (C) Sensorgram showing the immobilisation of dsDNA on the SA chip surface (association: dark grey phase; immobilised dsDNA: light grey phase). (D) Following dsDNA immobilisation, buffer containing 1 μM RecA(E38K) (+ 0.5 mM ATPγS) was flowed into the flow cell, at t = 0 min for 500 s. During this period, RecA(E38K) associated with ssDNA (blue phase), forming a RecA(E38K) filament. From 500 – 3,000 s, buffer containing 0.5 or 1 mM ATPγS was flowed into the flow cell (yellow phase). From 3,000 – 7,200 s, buffer containing 1 μM RecA(E38K) (+ 0.5 mM ATPγS) was flowed into the flow cell to allow for more RecA(E38K) to associate with the dsDNA. (E) Sensorgram showing the association of pol IV with RecA(E38K) structures formed on dsDNA. At t = 0 s, 0.65 uM pol IV (+ 0.5 mM ATPγS) was flowed into the flow cell for 220 s and association of pol IV was observed (green phase). At t = 220 s, buffer containing 0.5 mM ATPγS was flowed into the flow cell (yellow phase). (F) Sensorgram showing the association of pol IV with dsDNA. At t = 0 s, 0.65 uM pol IV (+ 0.5 mM ATPγS) was flowed into the flow cell for 220 s and association of pol IV was observed (green phase). At t = 220 s, buffer containing 0.5 mM ATPγS was flowed into the flow cell (yellow phase). Lower response units are recorded than for the association of pol IV with RecA(E38K) structures on dsDNA.

**Supplementary Figure S4. RecA(E38K) forms RecA*-like structures on circular dsDNA.** (A) RecA(E38K) readily binds to dsDNA. In six separate reactions, either RecA(E38K) or wild-type RecA was incubated at 37°C with nicked circular dsDNA (cdsDNA), ATP, and an ATP regeneration system. (B) LexA Cleavage Assays. Reaction mixtures contained 40 mM Tris-HCl at pH 8.0, 10 mM MgCl_2_, 30 mM NaCl, 2 mM dithiothreitol, 3 µM circular single-stranded DNA (cssDNA) or nicked circular double-stranded DNA (cdsDNA), 3 mM ATPγS, LexA, and RecA as noted. Reactions were incubated at 37°C for 10 minutes before addition of UmuD or LexA. The reaction products were separated and visualized by 15% SDS-PAGE stained with Coomassie blue. Lane 1 contains a protein ladder while subsequent groups of three lanes contain the same reaction mixture sampled at 0, 20, and 40 minutes. On cssDNA, RecA(E38K) and wild-type RecA form RecA* structures. On cdsDNA however, RecA(E38K) forms RecA*-like structures in contrast to wildtype RecA.

## Supplementary Notes

Sequence of pJMuvrA-PAmCherry-mCI vector:

AAGCTGGAAGATCTTCCCTGGCACGACAGGTTTCCCGACTGGAAAGCGGGCAGTGAGCGCAACGCAATTAATGTGAGTTAGCTCACTCATTAGGCACCCCAGGCTTTACACTTTATGCTTCCGGCTCGTATGTTGTGTGGAATTGTGAGCGGATAACAATTTCACACAGGAAACAGCTATGACCATGATTACGCCAAGCGCGCAATTAACCCTCACTAAAGGGAACAAAAGCTGGGTACCGGGCCCCCCCTCGAGGTCGACTTCCGGGAAACAAACCTGGCCAGACATTGTTACACAACACTCCGGGTAATGCATTCCAATACTGTATATTCATTCAGGTCAATTTGTGTCATAATTAACCGTTTGTGATCGGATCCAGCACCATGCCACCGGGCAAAAAAG CGTTTAATCCGGGAAAGCAT<u>ATGGTGAGCAAGGGCGAGGAGGATAACATGGCCATCATTAAGGAGTTCATGCGCTTCAAGGTGCACATGGAGGGGTCCGTGAACGGCCACGTGTTCGAGATCGAGGGCGAGGGCGAGGGCCGCCCCTACGAGGGCACCCAGACCGCCAAGCTGAAGGTGACCAAGGGTGGCCCCCTGCCCTTCACCTGGGACATCCTGTCCCCTCAATTCATGTACGGCTCCAATGCCTACGTGAAGCACCCCGCCGACATCCCCGACTACTTTAAGCTGTCCTTCCCCGAGGGCTTCAAGTGGGAGCGCGTGATGAAATTCGAGGACGGCGGCGTGGTGACCGTGACCCAGGACTCCTCCCTGCAAGACGGTGAGTTCATCTACAAGGTGAAGCTGCGCGGCACCAACTTCCCCTCCGACGGCCCCGTAATGCAGAAGAAGACCATGGGCTGGGAGGCCCTCTCCGAGCGGATGTACCCCGAGGACGGCGCCCTGAAGGGCGAGGTCAAGCCGCGCGTGAAGCTGAAGGACGGCGGCCACTACGACGCTGAGGTCAAGACCACCTACAAGGCCAAGAAGCCCGTGCAGCTGCCCGGCGCCTACAACGTCAACCGCAAGTTGGACATCACCTCACACAACGAGGACTACACCATCGTGGAACAGTACGAAC</u>GTGCCGAGGGCCGCCACTCCACCGGCGGCATGGACGAGCTGTACAAGGAGCTCGCT<u>GCAGGTGGCGGCGGCGGCTCCGGCAGCCATATGTATGAGTACCCTGTTTTTTCTCATGTTCAGGCAGGGATGTTCTCACCTGAGCTTCGCACCTTTACCAAAGGTGATGCGGAGCGCTGGGTAAGCACAACCAAAAAAGCCAGTGATTCTGCATTCTGGCTTGAGGTTGAAGGTAATTCCATGACCACACCAACAGGCTCCAAGACAAGCTTTCCTGACGGAATGTTAATTCTCGTTGACCCTGAGCAGGCTGTTGAGCCAGGTGATTTCTGCATTGCCCGCCTTGGGGGTGATGAGTTTACCTTCGCGAAACTGATCCGCGATAGCGGTCAGGTGTTTTTACAACCACTGAACCCACAGTACCCAATGATCCCATGCAATGAGAGTTGTTCCGTTGTGGGGAAAGTTATCGCTAGTCAGTGA</u>GCGGCCGCGAATTCGAAGTTCCTATAGTTTCTAGAGAATAGGAACTTCGATCTTTAGAAAAACTCATCGAGCATCAAATGAAACTGCAATTTATTC ATATCAGGATTATCAATACCATATTTTTGAAAAAGCCGTTTCTGTAATGAAGGAGAAAACTCACCGAGGCAGTTCCATAGGATGGCAAGATCCTGGTATCGGTCTGCGATTCCGACTCGTCCAACATCAATACAACCTATTAATTTCCCCTCGTCAAAAATAAGGTTATCAAGTGAGAAATCACCATGAGTGACGACTGAATCCGGTGAGAATGGCAAAAGCTTATGCATTTCTTTCCAGACTTGTTCAACAGGCCAGCCATTACGCTCGTCATCAAAATCACTCGCATCAACCAAACCGTTATTCATTCGTGATTGCGCCTGAGCGAGACGAAATACACGATCGCTGTTAAAAGGACAATTACAAACAGGAATCGAATGCAACCGGCGCAGGAACACTGCCAGCG CATCAACAATATTTTCACCTGAATCAGGATATTCTTCTAATACCTGGAATGCTGTTTTCCCGGGGATCGCAGTGGTGAGTAACCATGCATCATCAGGAGTACGGATAAAATGCTTGAT GGTCGGAAGAGGCATAAATTCCGTCAGCCAGTTTAGTCTGACCATCTCATCTGTAACATCATTGGCAACGCTACCTTTGCCATGTTTCAGAAACAACTCTGGCGCATCGGGCTTCCCATACAATCGATAGATTGTCGCACCTGATTGCCCGACATTATCGCGAGCCCATTTATACCCATATAAATCAGCATCCATGTTGGAATTTAATCGCGGGCGCGAGCAAGACGTTTCCCGTTGAATATGGCTCATAACACCCCTTGTATTACTGTTTATGTAAGCAGACAGTTTTATTGTTCATGATGATATATTTTTATCTTGTGCAATGTAACATCAGAGATTTTGAGACACAACGTGGCTTTCCCCGCCCGCCCGATCCCCGGGTACCGAGCTCGAATTTCGACCAATTCGAAGTTCCTATACTTTCTAGAGAATAGGAACTTCCCGCGGTGGAGCTCCAATTCGCCCTATAGTGAGTCGTATTACGCGCGCTCACTGGCCGTCGTTTTACAACGTCGTGACTGGGAAAACCCTGGCGTTACCCAACTTAATCGCCTTGCAGCACATCCCCCTTTCGCCAGCTGGCGTAATAGCGAAGAGGCCCGCACCGATCGCCCTTCCCAACAGTTGCGCAGCCTGAATGGCGAATGGGACGCGCCCTGTAGCGGCGCATTAAGCGCGGCGGGTGTGGTGGTTACGCGCAGCGTGACCGCTACACTTGCCAGCGCCCTAGCGCCCGCTCCTTTCGCTTTCTTCCCTTCCTTTCTCGCCACGTTCGCCGGAAGATCTTCCAATTCCCGACAGTAAGACGGGTAAGCCTGTTGATGATACCGCTGCCTTACTGGGTGCATTAGCCAGTCTGAATGACCTGTCACGGGATAATCCGAAGTGGTCAGACTGGAAAATCAGAGGGCAGGAACTGCTGAACAGCAAAAAGTCAGATAGCACCACATAGCAGACCCGCCATAAAACGCCCTGAGAAGCCCGTGACGGGCTTTTCTTGTATTATGGGTAGTTTCCTTGCATGAATCCATAAAAGGCGCCTGTAGTGCCATTTACCCCCATTCACTGCCAGAGCCGTGAGCGCAGCGAACTGAATGTCACGAAAAAGACAGCGACTCAGGTGCCTGATGGTCGGAGACAAAAGGAATATTCAGCGATTTGCCCGAGCTTGCGAGGGTGCTACTTAAGCCTTTAGGGTTTTAAGGTCTGTTTTGT AGAGGAGCAAACAGCGTTTGCGACATCCTTTTGTAATACTGCGGAACTGACTAAAGTAGTGAGTTATACACAGGGCTGGGATCTATTCTTTTTATCTTTTTTTATTCTTTCTTTATTC TATAAATTATAACCACTTGAATATAAACAAAAAAAACACACAAAGGTCTAGCGGAATTTACAGAGGGTCTAGCAGAATTTACAAGTTTTCCAGCAAAGGTCTAGCAGAATTTACAGATACCCACAACTCAAAGGAAAAGGACTAGTAATTATCATTGACTAGCCCATCTCAATTGGTATAGTGATTAAAATCACCTAGACCAATTGAGATGTATGTCTGAATTAGTTGTTTTCAAAGCAAATGAACTAGCGATTAGTCGCTATGACTTAACGGAGCATGAAACCAAGCTAATTTTATGCTGTGTGGCACTACTCAACCCCACGATTGAAAACCCTACAAGGAAAGAACGGACGGTATCGTTCACTTATAACCAATACGCTCAGATGATGAACATCAGTAGGGAAAATGCTTATGGTGTATTAGCTAAAGCAACCAGAGAGCTGATGACGAGAACTGTGGAAATCAGGAATCCTTTGGTTAAAGGCTTTGAGATTTTCCAGTGGACAAACTATGCCAAGTTCTCAAGCGAAAAATTAGAATTAGTTTTTAGTGAAGAGATATTGCCTTATCTTTTCCAGTTAAAAAAATTCATAAAATATAATCTGGAACATGTTAAGTCTTTTGAAAACAAATACTCTATGAGGATTTATGAGTGGTTATTAAAAGAACTAACACAAAAGAAAACTCACAAGGCAAATATAGAGATTAGCCTTGATGAATTTAAGTTCATGTTAATGCTTGAAAATAACTACCATGAGTTTAAAAGGCTTAACCAATGGGTTTTGAAACCAATAAGTAAAGATTTAAACACTTACAGCAATATGAAATTGGTGGTTGATAAGCGAGGCCGCCCGACTGATACGTTGATTTTCCAAGTTGAACTAGATAGACAAATGGATCTCGTAACCGAACTTGAGAACAACCAGATAAAAATGAATGGTGACAAAATACCAACAACCATTACATCAGATTCCTACCTACATAACGGACTAAGAAAAACACTACACGATGCTTTAACTGCAAAAATTCAGCTCACCAGTTTTGAGGCAAAATTTTTGAGTGACATGCAAAGTAAGTATGATCTCAATGGTTCGTTCTCATGGCTCACGCAAAAACAACGAACCACACTAGAGAACATACTGGCTAAATACGGAAGGATCTGAGGTTCTTATGGCTCTTGTATCTATCAGTGAAGCATCAAGACTAACAAACAA AAGTAGAACAACTGTTCACCGTTACATATCAAAGGGAAAACTGTCCATATATGCACAGATGAAAACGGTGTAAAAAAGATAGATACATCAGAGCTTTTACGAGTTTTTGGTGCATT CAAAGCTGTTCACCATGAACAGATCGACAATGTAACAGATGAACAGCATGTAACACCTAATAGAACAGGTGAAACCAGTAAAACAAAGCAACTAGAACATGAAATTGAACACCTGAGACAACTTGTTACAGCTCAACAGTCACACATAGACAGCCTGAAACAGGCGATGCTGCTTATCGAATCAAAGCTGCCGACAACACGGGAGCCAGTGACGCCTCCCGTGGGGAAAAAATCATGGCAATTCTGGAAGAAATAGCGCTTTCAGCCGGCAAACCTGAAGCCGGATCTGCGATTCTGATAACAAACTAGCAACACCAGAACAGCCCGTTTGCGGGCAGCAAAACCCGTGGGAATTAATTCCCCTGCTCGCGCAGGCTGGGTGCCAAGCTCTCGGGTAACATCAAGGCCCGATCCTTGGAGCCCTTGCCCTCCCGCACGATGATCGTGCCGTGATCGAAATCCAGATCCTTGACCCGCAGTTGCAAACCCTCACTGATCCGCATGCCCGTTCCATACAGAAGCTGGGCGAACAAACGATGCTCGCCTTCCAGAAAACCGAGGATGCGAACCACTTCATCCGGGGTCAGCACCACCGGCAAGCGCCGCGACGGCCGAGGTCTTCCGATCTCCTGAAGCCAGGGCAGATCCGTGCACAGCACCTTGCCGTAGAAGAACAGCAAGGCCGCCAATGCCTGACGATGCGTGGAGACCGAAACCTTGCGCTCGTTCGCCAGCCAGGACAGAAATGCCTCGACTTCGCTGCTGCCCAAGGTTGCCGGGTGACGCACACCGTGGAAACGGATGAAGGCACGAACCCAGTGGACATAAGCCTGTTCGGTTCGTAAGCTGTAATGCAAGTAGCGTATGCGCTCACGCAACTGGTCCAGAACCTTGACCGAACGCAGCGGTGGTAACGGCGCAGTGGCGGTTTTCATGGCTTGTTATGACTGTTTTTTTGGGGTACAGTCTATGCCTCGGGCATCCAAGCAGCAAGCGCGTTACGCCGTGGGTCGATGTTTGATGTTATGGAGCAGCAACGATGTTACGCAGCAGGGCAGTCGCCCTAAAACAAAGTTAAACATCATGAGGGAAGCGGTGATCGCCGAAGTATCGACTCAACTATCAGAGGTAGTTGGCGTCATCGAGCGCCATCTCGAACCGACGTTGCTGGCCGTACATTTGTACGGCTCCGCAGTGGAT GGCGGCCTGAAGCCACACAGTGATATTGATTTGCTGGTTACGGTGACCGTAAGGCTTGATGAAACAACGCGGCGAGCTTTGATCAACGACCTTTTGGAAACTTCGGCTTCCCCTG GAGAGAGCGAGATTCTCCGCGCTGTAGAAGTCACCATTGTTGTGCACGACGACATCATTCCGTGGCGTTATCCAGCTAAGCGCGAACTGCAATTTGGAGAATGGCAGCGCAATGACATTCTTGCAGGTATCTTCGAGCCAGCCACGATCGACATTGATCTGGCTATCTTGCTGACAAAAGCAAGAGAACATAGCGTTGCCTTGGTAGGTCCAGCGGCGGAGGAACTCTTTGATCCGGTTCCTGAACAGGATCTATTTGAGGCGCTAAATGAAACCTTAACGCTATGGAACTCGCCGCCCGACTGGGCTGGCGATGAGCGAAATGTAGTGCTTACGTTGTCCCGCATTTGGTACAGCGCAGTAACCGGCAAAATCGCGCCGAAGGATGTCGCTGCCGACTGGGCAATGGAGCGCCTGCCGGCCCAGTATCAGCCCGTCATACTTGAAGCTAGACAGGCTTATCTTGGACAAGAAGAAGATCGCTTGGCCTCGCGCGCAGATCAGTTGGAAGAATTTGTCCACTACGTGAAAGGCGAGATCACCAAGGTAGTCGGCAAATAATGTCTAACAATTCGTTCAAGCCGACGCCGCTTCGCGGCGCGGCTTAACTCAAGCGTTAGATGCACTAAGCACATAATTGCTCACAGCCAAACTATCAGGTCAAGTCTGCTTTTATTATTTTTAAGCGTGCATAATAAGCCCTACACAAATTGGGAGATATATCATGAAAGGCTGGCTTTTTCTTGTTATCGCAATAGTTGGCGAAGTAATCGCAACATCCGCATTAAAATCTAGCGAGGGCTTTA CT

