## Supplementary Figure S1 for "Modulation of DNA polymerase IV activity by UmuD and RecA* observed by single-molecule time-lapse microscopy"

### Supplementary Figure 1

**A** **Colocalisation with 50 ms exposure:**
in *umuDC* mutants

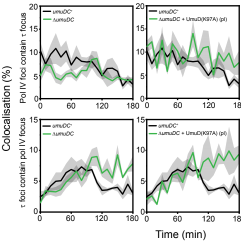


**B** **Foci per cell with 50 ms exposures:**
in *umuDC* mutants

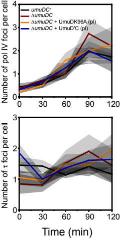
