## Supplementary figures and images for "Modulation of DNA polymerase IV activity by UmuD and RecA* observed by single-molecule time-lapse microscopy"

### Supplementary Figure S2

#### Supplementary Figure 2

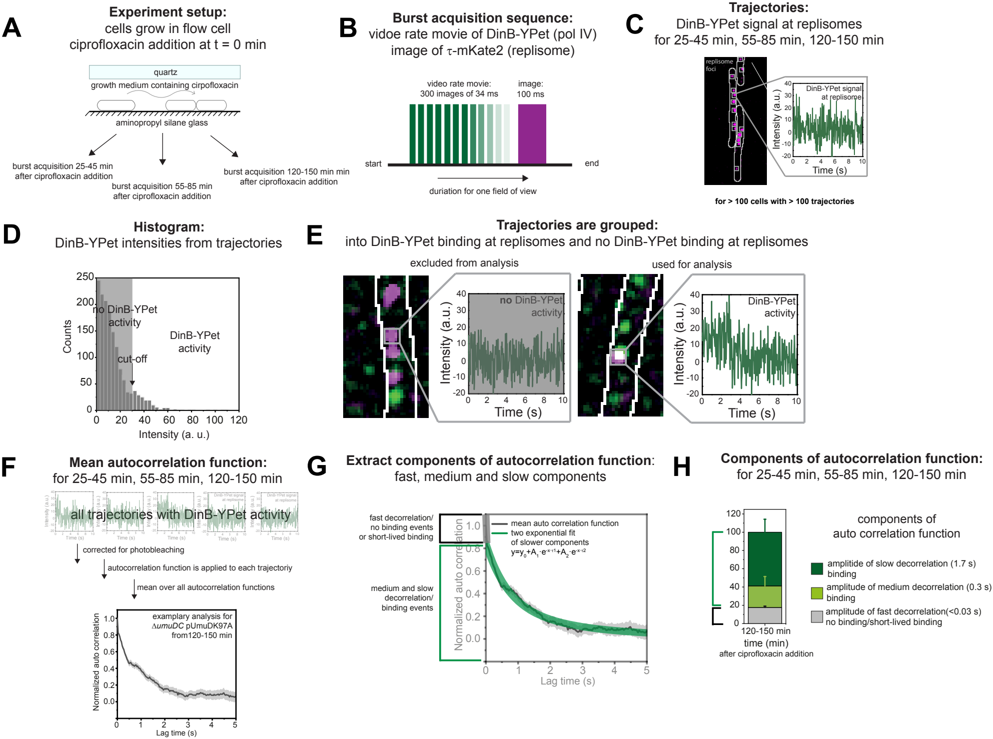

### Supplementary Figure S3

#### Supplementary Figure 3

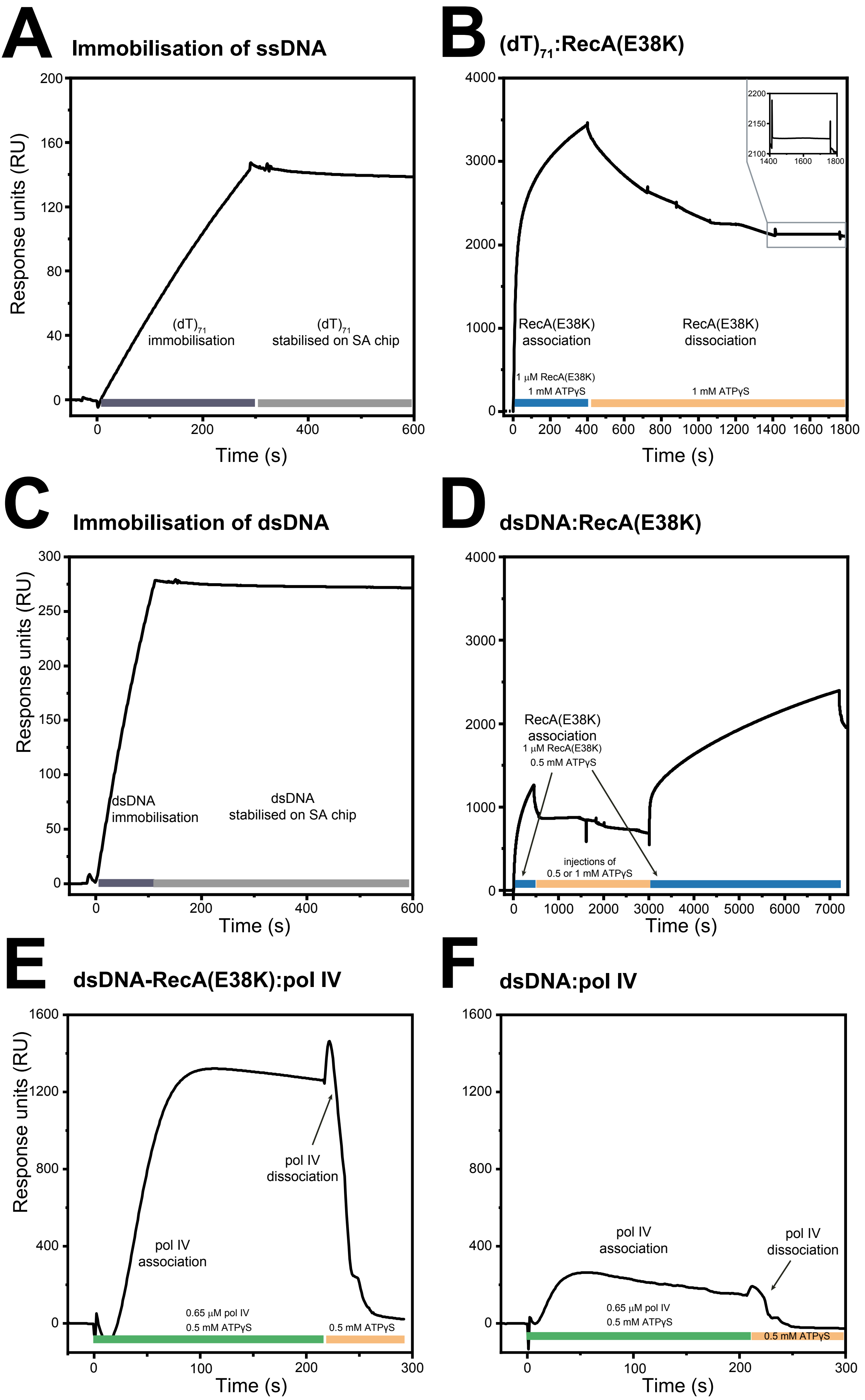

### Supplementary Figure S4

#### Supplementary Figure 4

**A** **ATPase hydrolysis assay**

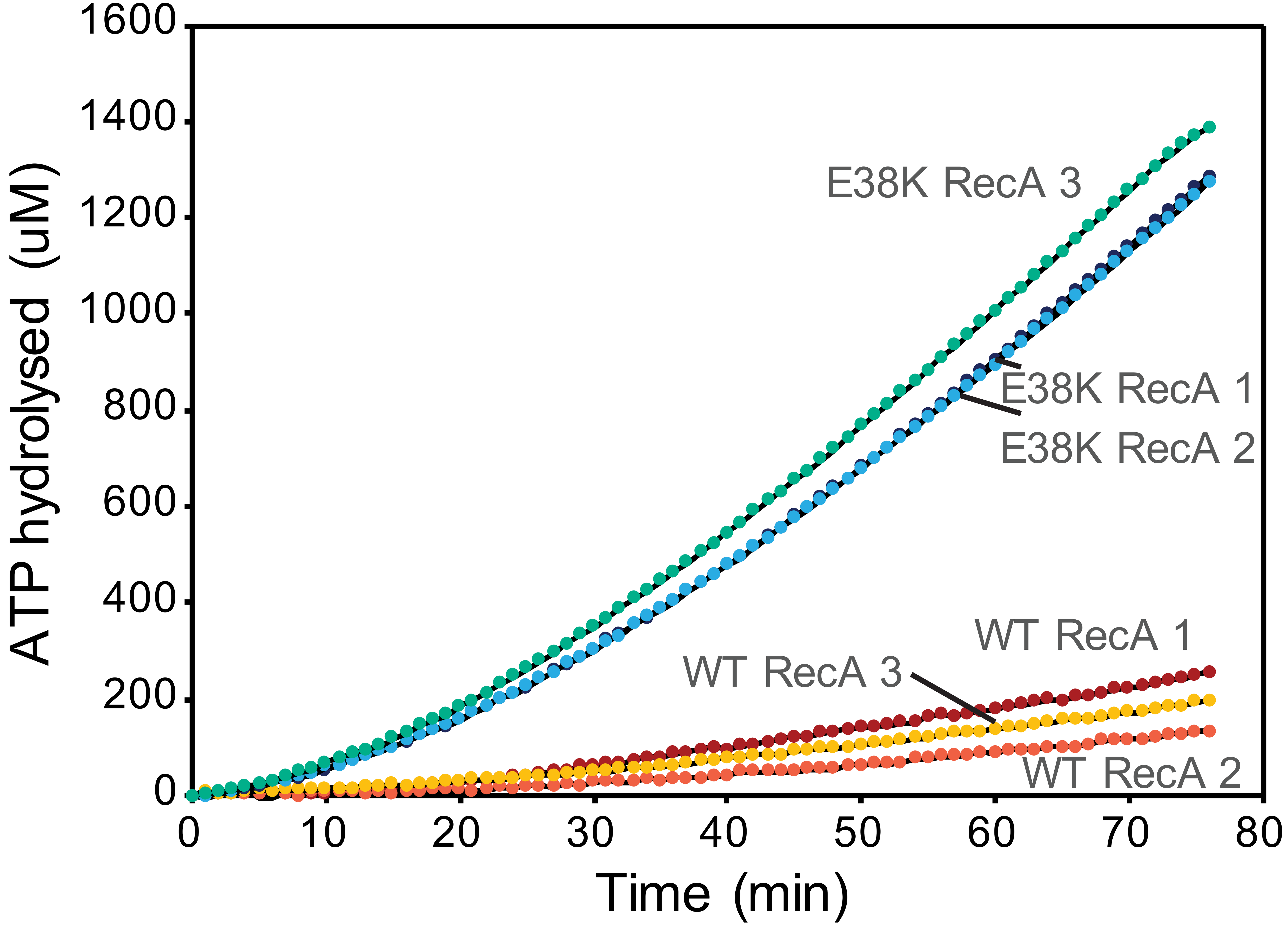


**B** **LexA Cleavage assay**

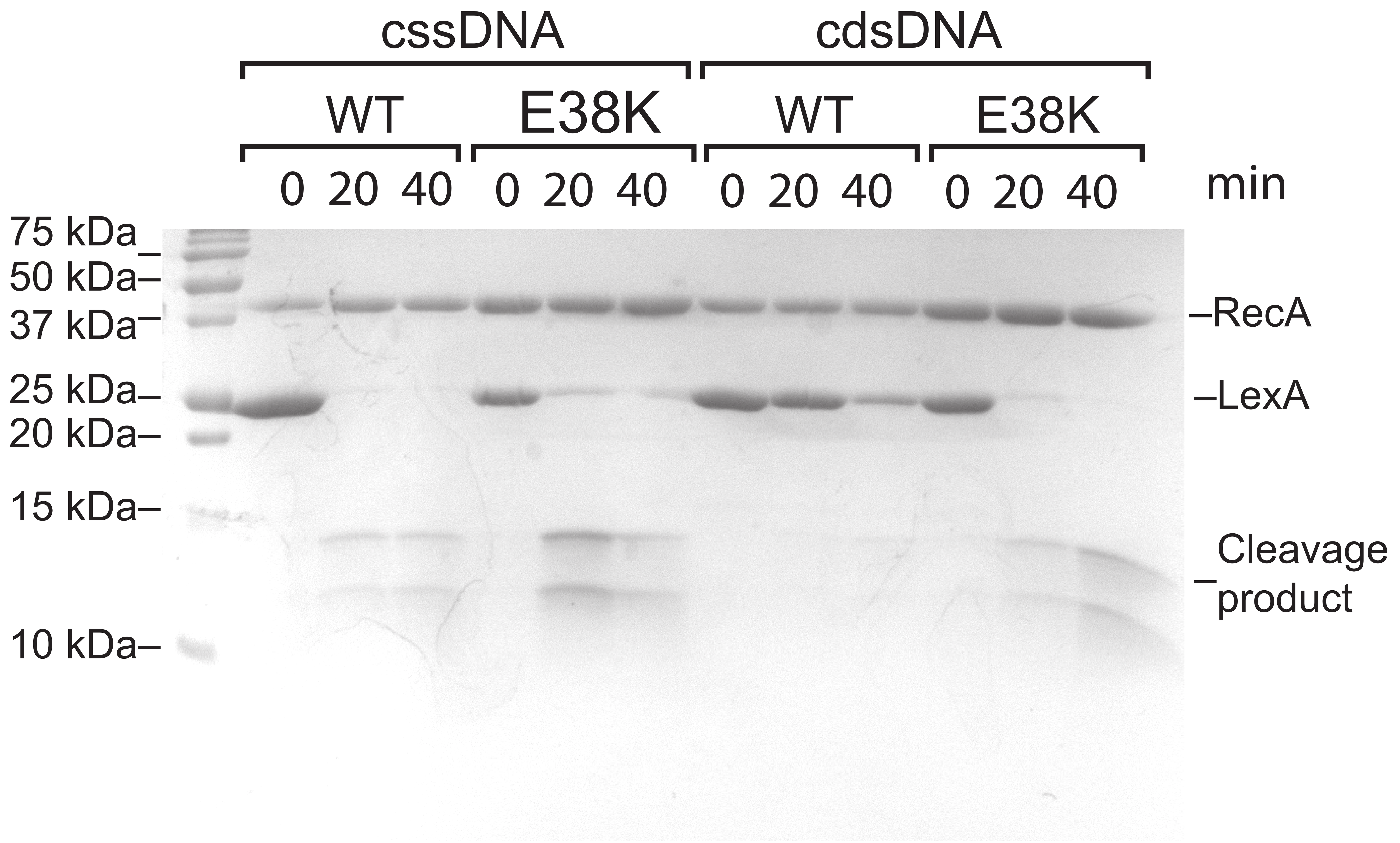
